# Skeletons in the closet: The importance of actin in alphavirus replication

**DOI:** 10.64898/2026.04.20.719692

**Authors:** Alyssa Zaye Lara, Richard W. Hardy, Melissa Phelps, Irene Newton

## Abstract

The ability of the bacterial endosymbiont *Wolbachia pipientis* to block arboviruses in its mosquito host may be impinged by host genetic variation, leading to reduced efficacy in field releases. Across a large collection of *Drosophila* lines carrying natural genetic variation, we found that viral replication varied greatly in the absence of *Wolbachia*. However, the introduction of the symbiont reduced viral load in each background to similar levels, near the limit of detection. Therefore, *Wolbachia*-mediated viral blocking is seemingly robust against host genetic background. A genome-wide association study harnessing the variation in the viral loads across the *Wolbachia*-free set identified *rhoGAP18B* and *betaCOP* as host factors that contribute to SINV replication; furthermore, the gene products of which seemingly interact with each other in the context of cytoskeletal dynamics. These results shed light on host requirements for SINV replication and suggest possible avenues by which *Wolbachia* may encroach upon them during blocking.

## INTRODUCTION

Arthropod-borne viruses (arboviruses)—particularly those transmitted by mosquitoes—pose a great public health risk to humans, as their pathology is severe, including encephalitis, hemorrhaging, arthralgia, and death^1,2^. These viruses typically have RNA as genomic material, and can come from the Flaviviridae, Togaviridae, Reoviridae, and Bunyaviridae families^3^. There are currently no effective vaccines or treatments available^1^, and the rise in global warming is causing the mosquito vectors to invade habitats where they are not typically found^4^. Conventional arboviral prevention strategies revolve around limiting host propagation, such as through fumigation. However, this is not ideal since insecticides are non-discriminant—killing non-target insects and leading to an increase in insecticide resistance^5^. Others have attempted to genetically engineer mosquitoes to be recalcitrant to arboviruses; for example, they have generated mosquitoes that produce RNAi targeting arboviral genomes^6^, or mosquitoes that code for an antibody to neutralize arboviruses^7^. While promising, releasing such genetically modified organisms into the wild is not supported by many communities, and government regulations make this strategy infeasible in many locations. Hence, better strategies to combat arboviruses are needed.

A promising candidate for vector control is the Gram-negative, endosymbiotic bacterium *Wolbachia pipientis*. It naturally colonizes ∼40-52% of insect species^8^, and can block the replication of arboviruses in mosquitoes, rendering them incapable of arboviral transmission^9,10^—this phenomenon is termed “viral blocking”. While *Wolbachia* has not been found to naturally infect *Aedes aegypti* (the principal vector of major arboviruses such as dengue (DENV), chikungunya (CHIKV), and Zika (ZIKV)), the mosquito has been successfully transinfected with *Wolbachia* from *Drosophila melanogaster* (*Wolbachia* strain *w*Mel)^11^. The efficacy of *Wolbachia* as an arboviral prevention strategy has been realized, as the World Mosquito Program has been actively releasing *w*Mel-transinfected *Ae. aegypti* into endemic areas, and this has resulted in a decrease in dengue cases^12^.

Despite the success of the *Wolbachia* strategy, a few studies have suggested that host genetic background could influence the efficacy of viral blocking. Specifically, experimental evolution for weak or strong blocking mosquito strains identified variation in the host associated with a change in phenotype^13^, while naturally-occurring host genetic variation across two mosquito populations has been shown to influence the degree of viral blocking^14^. Both studies suggest a potential pitfall to the use of *Wolbachia* in limiting viral replication, prompting the question: *how robust is pathogen blocking to host genetic variation*?

To this end, the *Drosophila melanogaster* Genetic Reference Panel—a large collection of genetically distinct fly lines harboring naturally occurring variations—was used to identify potential variation in viral blocking^15,16^. The fly lines have each been sequenced, and a comparison of their response to virus in the presence and absence of the symbiont permits the evaluation of the impact of *Wolbachia* on variation in viral replication across the DGRP. This study demonstrates that *Wolbachia*-mediated viral blocking is surprisingly robust to host genetic background, but that SINV interacts with the host cytoskeleton and endocytic machinery during infection, while *Wolbachia* restructures both the actin and microtubule cytoskeletons. It is likely that the robust changes which *Wolbachia* induces in its host’s cells during infection eclipse any influence of genetic variation observed in the DGRP.

## RESULTS

### Host genetic diversity does not alter viral blocking

The *Drosophila melanogaster* Genetic Reference Panel (DGRP) is a collection of 205 fly lines originally obtained from North Carolina, then inbred for more than 20 generations^15,16^. All lines have been fully sequenced, and the panel has been used by many researchers to investigate various phenotypes^17,18^. Since these lines were originally wild-type, the genetic variation present across the set is naturally-occurring; furthermore, the large number of fly lines makes this panel a powerful tool to use in this screen. Of the 205 lines, those that harbor *Wolbachia* were selected for use in this study; these were then treated with tetracycline to generate a *Wolbachia*-negative set. Both the *Wolbachia*-positive and *Wolbachia*-negative sets were injected with Sindbis virus tagged with nanoluciferase on the nsP3 gene (SINV-nsP3-nLuc). Using this tagged virus, the resulting viral loads can be measured using a luciferase assay. For each of the DGRP lines, five *Wolbachia*-positive and five *Wolbachia*-negative individuals were injected and analyzed (**Figure 1**).

**Figure 1.**
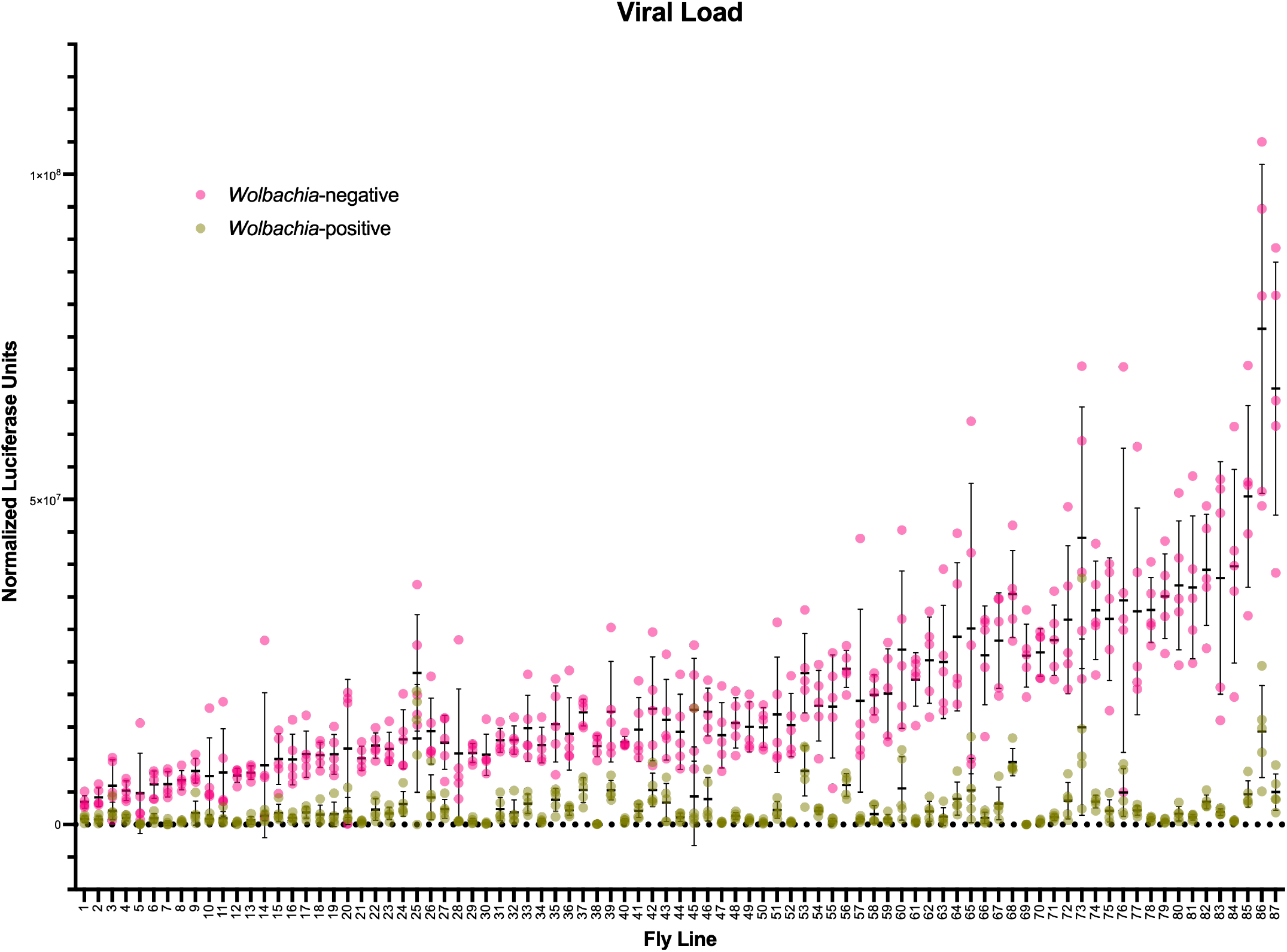
*Wolbachia*-mediated viral blocking is seemingly robust to host genetic variation, while SINV replication is heavily influenced by it. The *Wolbachia*-positive lines of the DGRP, as well as their tetracycline-cleared counterparts, were injected with Sindbis virus, and the resulting viral loads measured. The luciferase values serve as a proxy for viral titer. Each data point is the viral load of a single fly (*n*=5 for *Wolbachia*-positive, *n*=5 for *Wolbachia*-negative). Each line of the DGRP is genetically distinct. In the absence of *Wolbachia* (pink), viral load varies greatly; meanwhile, in the presence of *Wolbachia* (green), viral load is hedged to a ceiling. Thus, SINV replication is sensitive to the natural genetic variation exhibited by the host.

The resulting viral loads in the *Wolbachia*-positive set are consistently lower compared to the *Wolbachia*-negative set, displaying the characteristic effect of *Wolbachia*-mediated viral blocking. On average, the presence of *Wolbachia* restricts viral replication to one-eighth that of the *Wolbachia*-negative set. These fly lines are genetically distinct from one another, and therefore it is evident that viral replication itself is heavily affected by host genetic background since across the *Wolbachia*-negative set, the resulting viral loads have such a wide range—specifically, 23% of the fly lines have *Wolbachia*-negative viral loads with a |*z*-score| greater than 1. Interestingly, even in genetic backgrounds where viral replication is high, *Wolbachia* is still able to restrict the resulting viral load to a similarly low magnitude when it is present, as evidenced by the narrow range of viral values in the *Wolbachia*-positive set. In fact, across the *Wolbachia*-positive set, only 8% of the fly lines have viral loads with a *z*-score greater than 1. Thus, *Wolbachia* seemingly reduces viral replication to a ceiling, regardless of host genetic background.

### Blocking strength is not associated with *Wolbachia* density in the DGRP

Host genetic variation has been found to affect *Wolbachia* density, and *Wolbachia* density, in turn, has previously been correlated with blocking efficiency^19^. Thus, to identify whether there was any correlation between the magnitude of viral replication restricted by the presence of *Wolbachia* and *Wolbachia* titers in individual flies, the loads of *Wolbachia* harbored by the fly lines across the DGRP were measured by qPCR (*n*=10) using primers targeting *wsp* (*Wolbachia* surface protein), normalized to the housekeeping gene *rpL32* (large ribosomal subunit protein eL32). Across the DGRP, there is some variation in *Wolbachia* density (**Figure 2**a); specifically, 18% of the fly lines have a *Wolbachia* titer with a |*z*-score| greater than 1. However, a linear correlation analysis of *Wolbachia* density and blocking strength (defined as the difference between viral replication in the presence or absence of *Wolbachia*) failed to show a relationship between the two (R^2^ = 0.01000) (**Figure 2**b). Thus, while *Wolbachia* density may be impacted by the naturally-occurring genetic variation present across the DGRP, it does not correlate with viral blocking strength, as defined here.

**Figure 2.**
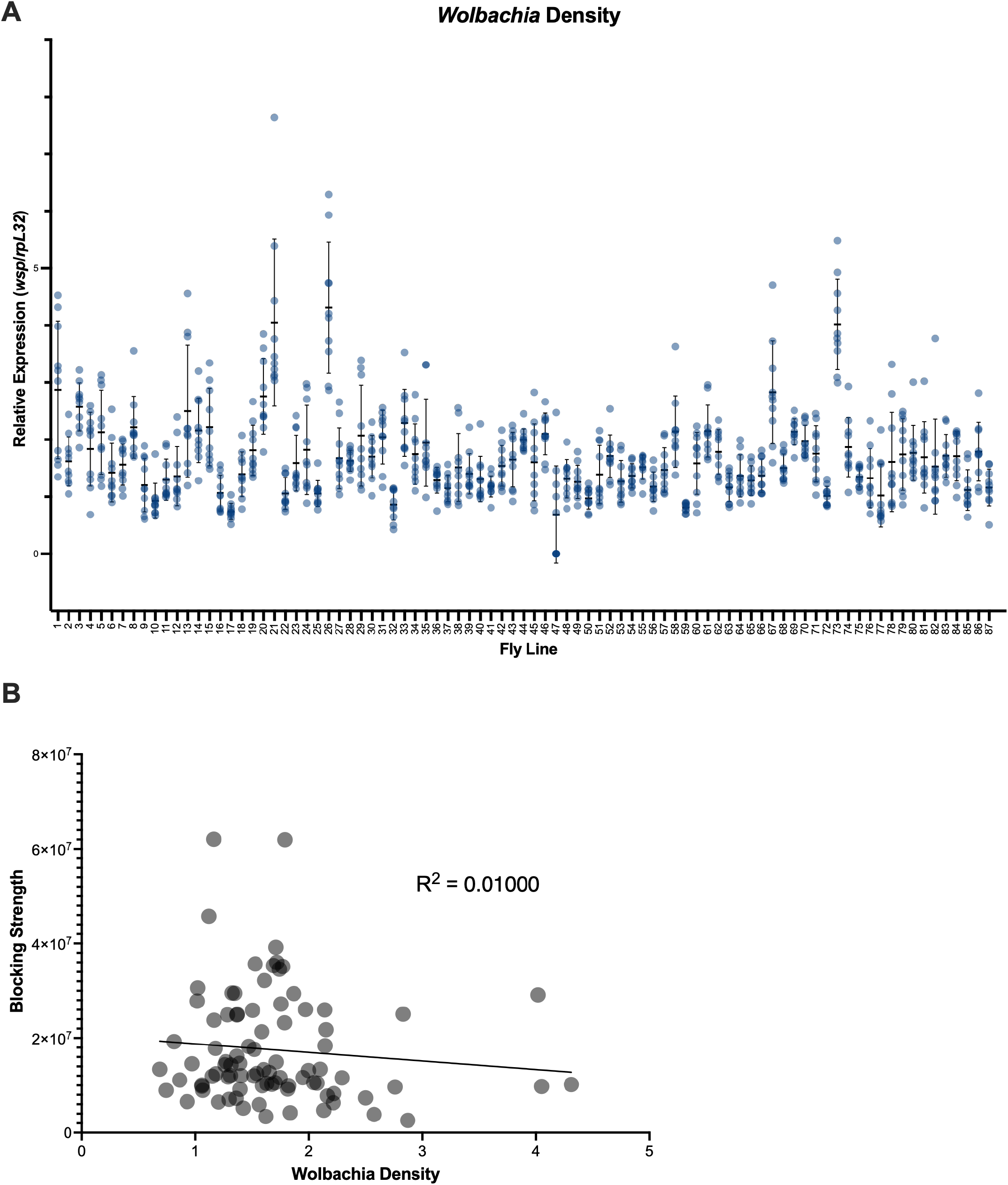
Blocking strength across the DGRP lines does not correlate with *Wolbachia* density. (A) Using qPCR, *Wolbachia* density is measured as the proportion of *Wolbachia* genome copies (using *wsp*) in reference to a host housekeeping gene (*rpL32*). Each data point is the *Wolbachia* density of a single fly (*n*=10). (B) Linear correlation analysis between *Wolbachia* density and viral blocking strength. The absence of a linear relationship indicates that blocking strength is not dependent on *Wolbachia* density in the DGRP.

### A genome-wide association study for SINV replication

Since the primary variation in the viral load dataset came from *Wolbachia*-free flies, we reasoned that host genetic determinants associated with SINV replication could be identified by running a genome-wide association study using the DGRPool web tool^20^. The viral load of each *Wolbachia*-positive line was subtracted from the viral load of its genetically identical *Wolbachia*-negative line and used as input. Each SNP in the DGRP, and its contribution to the extent of SINV replication in that fly background, was visualized using a Manhattan plot (**Figure 3**a), and emphasis was accorded to genes supported by numerous, highly significant SNPs. For example, the SNP with the most significant *p*-value (the tallest peak in the Manhattan plot) falls within the gene *pyd*; however, this gene holds a total of only 5 SNPs from the GWAS. Similarly, the SNP with the second-most significant *p*-value falls within the gene *pka-C2*, which holds a total of only 2 SNPs. The bubble plot in **Figure 3**b permits the visualization of gene regions that have many highly significant SNPs. Here, each annotated gene locus is plotted according to its most significant SNP, and the size of its plot point corresponds to how many highly significant SNPs fell within its gene region. **Table 1** lists the gene regions that contain the greatest number of highly significant SNPs from the GWAS.

**Figure 3.**
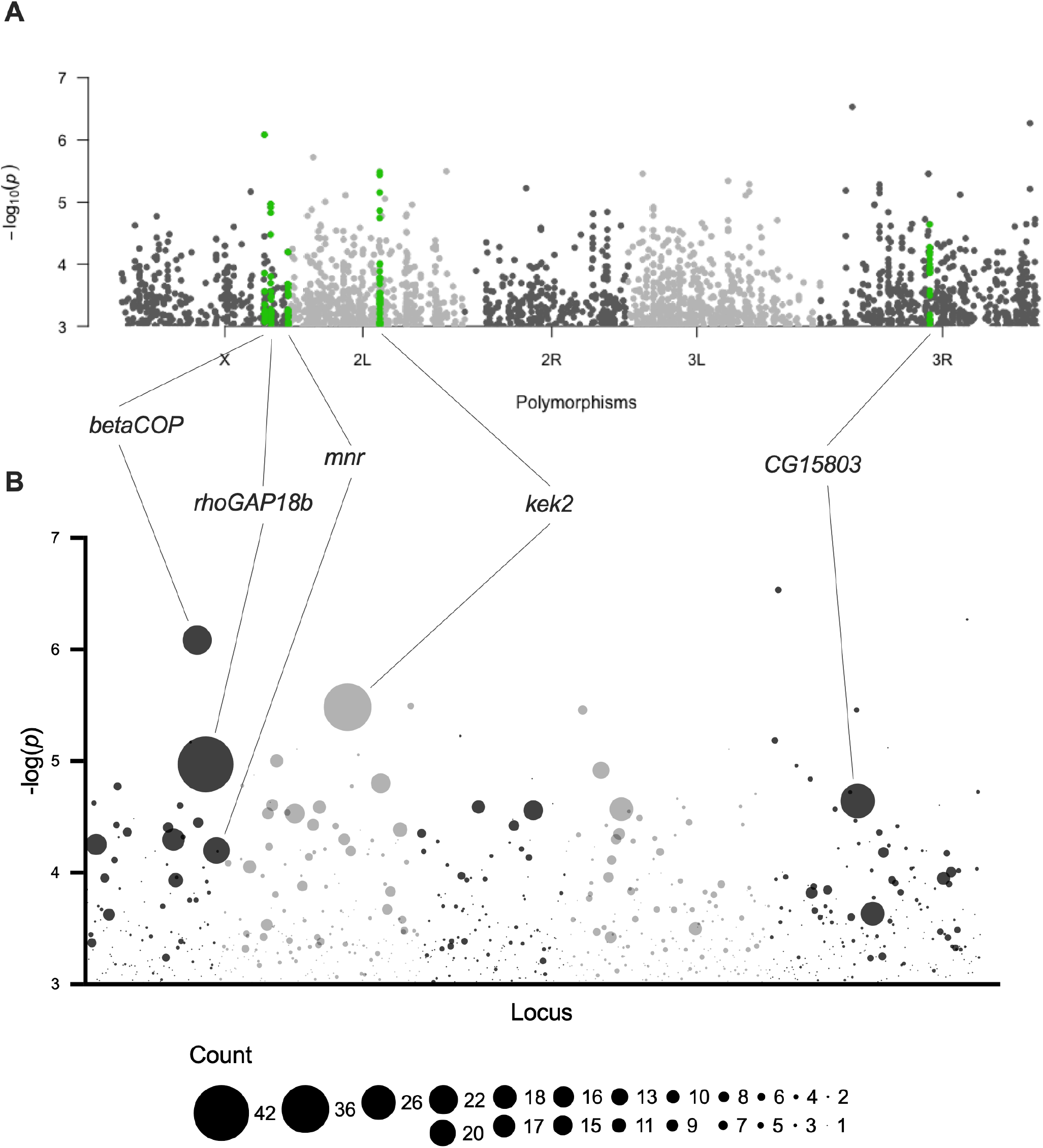
The actin cytoskeletal regulator *rhoGAP18B* is highly associated with SINV replication. (A) The GWAS analysis provided how significantly associated each SNP present in the DGRP is with SINV replication. In this Manhattan plot, each SNP is plotted to its corresponding significance value (*p*-value). The most significant SNPs were included in this graph (*p*-value < 1×10^-3^). (B) Several loci had a greater number of highly significant SNPs fall within their gene region, compared to others. In this bubble plot, each annotated gene locus is plotted according to its most significant SNP, and the size of its plot point corresponds to how many highly significant SNPs fell within its gene region.

**Table 1.** Top hits from the GWAS. Annotated genes with the greatest number of highly significant SNPs (*p*<1×10^-3^) within their gene region. *p*-value_max_ refers to the *p*-value of the SNP within the indicated gene region that was most significant (top point in the Manhattan plot peak).

| Chromosome | Gene | # of SNPs | $p\text{-value}_{\max}$ |
| --- | --- | --- | --- |
| X | *rhoGAP18B* | 42 | $1.07 \times 10^{-5}$ |
| 2L | *kek2* | 36 | $3.29 \times 10^{-6}$ |
| 3R | *CG15803* | 26 | $2.28 \times 10^{-5}$ |
| X | *betaCOP* | 22 | $8.25 \times 10^{-7}$ |
| X | *mnr* | 20 | $6.34 \times 10^{-5}$ |

Using PANGEA^21^, a gene set enrichment analysis was conducted on the annotated gene regions that contained highly significant SNPs (*p*-value cutoff: 1×10^-3^). The datasets included in the analysis were the *Drosophila* biological process gene ontology subset and the FlyBase signaling pathway database, and the enriched gene sets considered were those with *p*-value < 0.01. The biological process with the highest fold enrichment is <u>actin cytoskeletal organization</u> (**Supplementary Figure 1**a), consistent with the involvement of such as indicated by the top hits *rhoGAP18B* and *betaCOP* (**Table 1**). This is followed by <u>cell adhesion</u> and <u>cell morphogenesis</u>, both of which are intrinsically linked to actin dynamics^22^.

Our gene set enrichment results focused our project on *rhoGAP18B*, a regulator of actin organization^23,24^ which holds 42 highly significant SNPs within its gene region—the most in the GWAS data set. Meanwhile, 22 highly significant SNPs fall within the gene region of *betaCOP*, which is part of the adaptor subcomplex of COPI, the coat protein complex involved in ER-Golgi vesicular transport^25^ Interestingly, knockdown of *betaCOP* has been shown to result in decreased replication of another alphavirus, CHIKV^26^. Prior work connects these two proteins in vesicular transport as the COPI coatomer complex at the Golgi involves actin—specifically, the COPI coatomer recruits activated Cdc42 and Rac to the Golgi by direct interaction to gammaCOP (another member of the COPI complex); Cdc42 then assembles actin through the Arp2/3 complex^27,28^. Furthermore, *rhoGAP18B* functions to inactivate Cdc42 and Rac1^23^. Taken together, the top hits of the GWAS seem to indicate the importance of the actin cytoskeleton to SINV replication, although how these particular alleles alter the function of these proteins or the structure of the cytoskeleton is not known.

In addition, several signaling pathways—Notch, FGFR, EGFR, and Hippo—were also enriched in the PANGEA analysis (**Supplementary Figure 1**b). While these pathways are canonically studied for their contributions to development, cell differentiation, and proliferation^29–32^, they have also been associated with viral replication^33–37^. Interestingly, all four of these pathways are also involved in actin dynamics^38–45^.

In all, a GWAS for SINV replication indicates the importance of the genes *rhoGAP18B* and *betaCOP*, which are likely exerting influence through their involvement in cytoskeletal dynamics.

### Role of *rhoGAP18B* in SINV replication

*rhoGAP18B* is a member of a family of twenty-one GTPase activating proteins that regulate the Rho family of GTPases^46^. It has four main isoforms, designated A through D (**Figure 4**a). Isoforms A and B are almost completely overlapping, save for a small portion of the first exon of isoform B that is missing in isoform A. Similarly, isoforms C and D are almost completely overlapping, save for a portion of the first exon of isoform C that is missing in isoform D. All four isoforms contain the RhoGAP domain on the C-terminal end^47^. These isoforms have previously been shown to regulate different actin pathways that result in different cytoskeletal dynamics^23^. To examine the effect of *rhoGAP18B* on viral replication, the different isoforms can be knocked down, and the resulting viral replication measured. We used existing whole fly RNAi resources, targeting the different isoforms; **Figure 4**a indicates the regions targeted by these RNAi constructs (green boxes).

**Figure 4.**
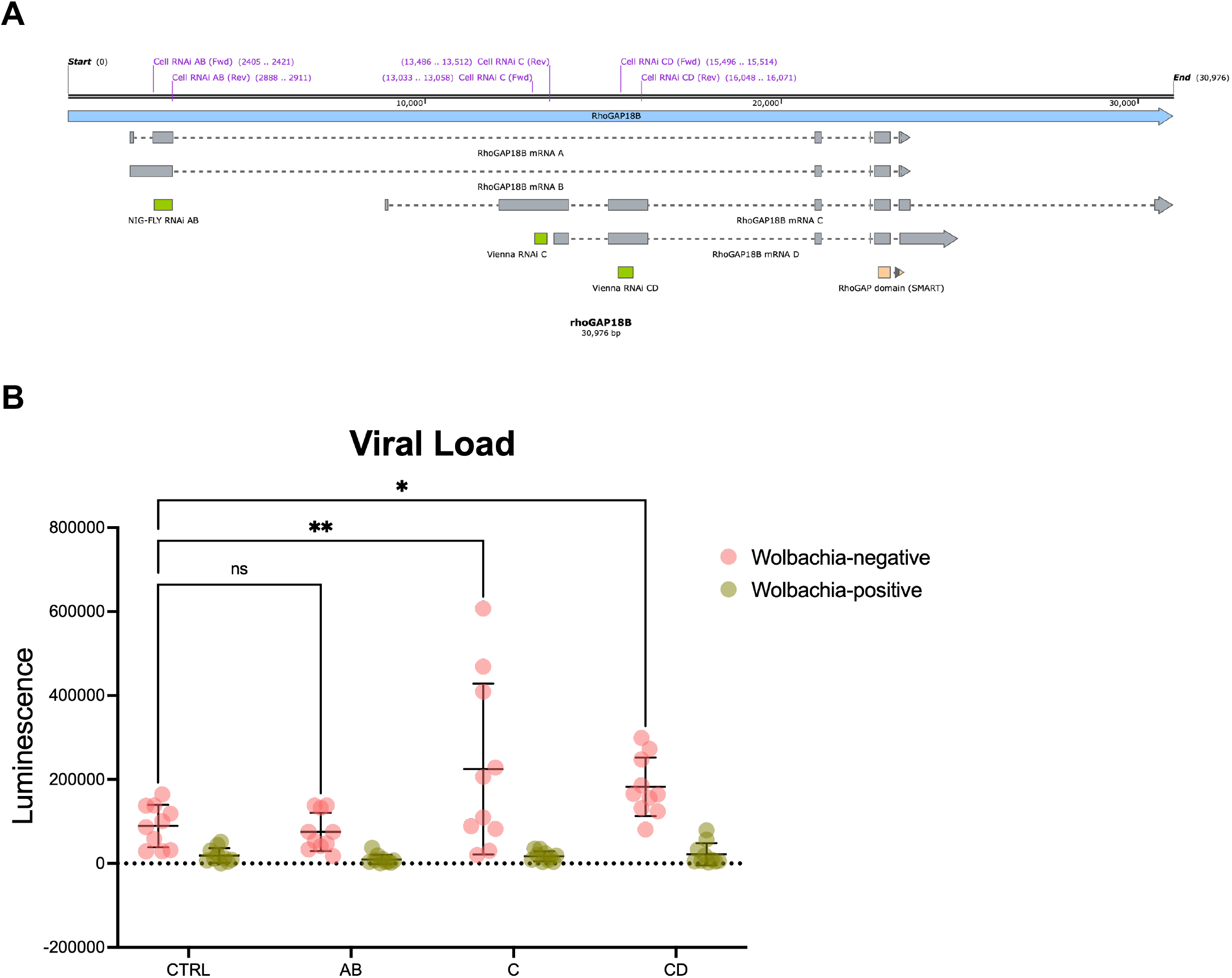
*rhoGAP18B* behaves in an anti-viral manner. (A) Gene structure of *rhoGAP18B. rhoGAP18B* has four isoforms, all of which terminate in the RhoGAP domain. The regions targeted by the RNAi constructs used in this study are indicated—constructs for whole fly in green, and for cell culture in purple. (B) Effect of *rhoGAP18B* knockdown on viral load. GAL4/UAS system was used to drive the expression of RNAi constructs targeting the different *rhoGAP18B* isoforms, resulting in knockdown of the indicated isoforms in the presence (green) and absence (pink) of *Wolbachia*. Flies were injected with Sindbis virus, and the resulting viral loads measured. The luciferase values serve as a proxy for viral titer. Each data point is the viral load of a single fly (*n*=10 for each condition). CTRL has the same genetic background as the experimental, but harbors an empty UAS vector. Two-way ANOVA; CTRL vs. A adjusted *p*-value=0.9722; CTRL vs. C adjusted *p*-value=0.0010; CTRL vs. CD adjusted *p*-value=0.0346.

In the whole fly, the RNAi constructs are under the control of the GAL4/UAS system. Thus, the three knockdown fly lines (isoforms AB, C, and CD), as well as the control—a fly line of similar genetic background (w^1118^) that contains the empty vector—were crossed to females from the Actin-GAL4 driver line, both with and without *Wolbachia*. The resulting progeny were then injected with SINV-nsP3-nLuc, and the resulting viral loads measured (**Figure 4**b). Across all the fly lines, viral replication in the *Wolbachia*-positive flies were consistently lower than in their *Wolbachia*-negative counterparts, indicating that even in the loss of *rhoGAP18B*, viral blocking still occurs. Notably, in *Wolbachia*-negative flies, loss of the C and CD isoforms resulted in increased viral replication (Two-way ANOVA; CTRL vs C-knockdown: *p*=0.0010; CTRL vs CD-knockdown: *p*=0.0346). Thus, these isoforms behave in an anti-viral manner.

### Cellular intersection of SINV and *Wolbachia*

Despite the genetic variation present across the entire DGRP set, viral blocking was consistently strong; variation in SINV replication was only observed when *Wolbachia* was absent. That said, there is striking overlap between the gene set enrichment analysis results and cellular pathways impacted by *Wolbachia*. For example, *Wolbachia* colonization is influenced by cell morphogenesis^48^, and the symbiont alters host actin^49,50^, endocytosis^51^, and vesicular trafficking^52^. Meanwhile, alphaviruses are known to induce morphological changes in the cell to produce structures that bolster their replication^53^; and cell adhesion has previously been associated with viral blocking^13,54,55^. Furthermore, pathway enrichment in screens for host genes associated with *Wolbachia* colonization^56,57^, viral infection^58^, and *Wolbachia* viral blocking^54,59^ have repeatedly indicated the involvement of cytoskeletal organization. We therefore reasoned that *Wolbachia* infection in and of itself may alter cytoskeletal dynamics in the cell. We used *Drosophila* cell lines, infected with the symbiont or not, to explore the morphology of the actin and microtubule cytoskeletons. Indeed, there are dramatic differences in the actin cytoskeleton and microtubule network when *Wolbachia* is present in the cell, the most prominent of which is the appearance of filopodia-like extensions (**Figure 5**a). Because viruses also hijack the host cytoskeleton^53,60^, it is possible that *Wolbachia*’s manipulation of this important cellular network might impinge upon viral replication. Because knockdown of *rhoGAP18B* results in morphological changes of the actin cytoskeleton^23^, we sought to confirm this result in our system. Interestingly, RNAi-mediated loss of *rhoGAP18B* in *Drosophila* cells resulted in a filopodia-like morphology as previously observed (**Figure 5**bc). Therefore, both the presence of *Wolbachia* and alterations to *rhoGAP18B* expression modify the host cytoskeleton.

**Figure 5.**
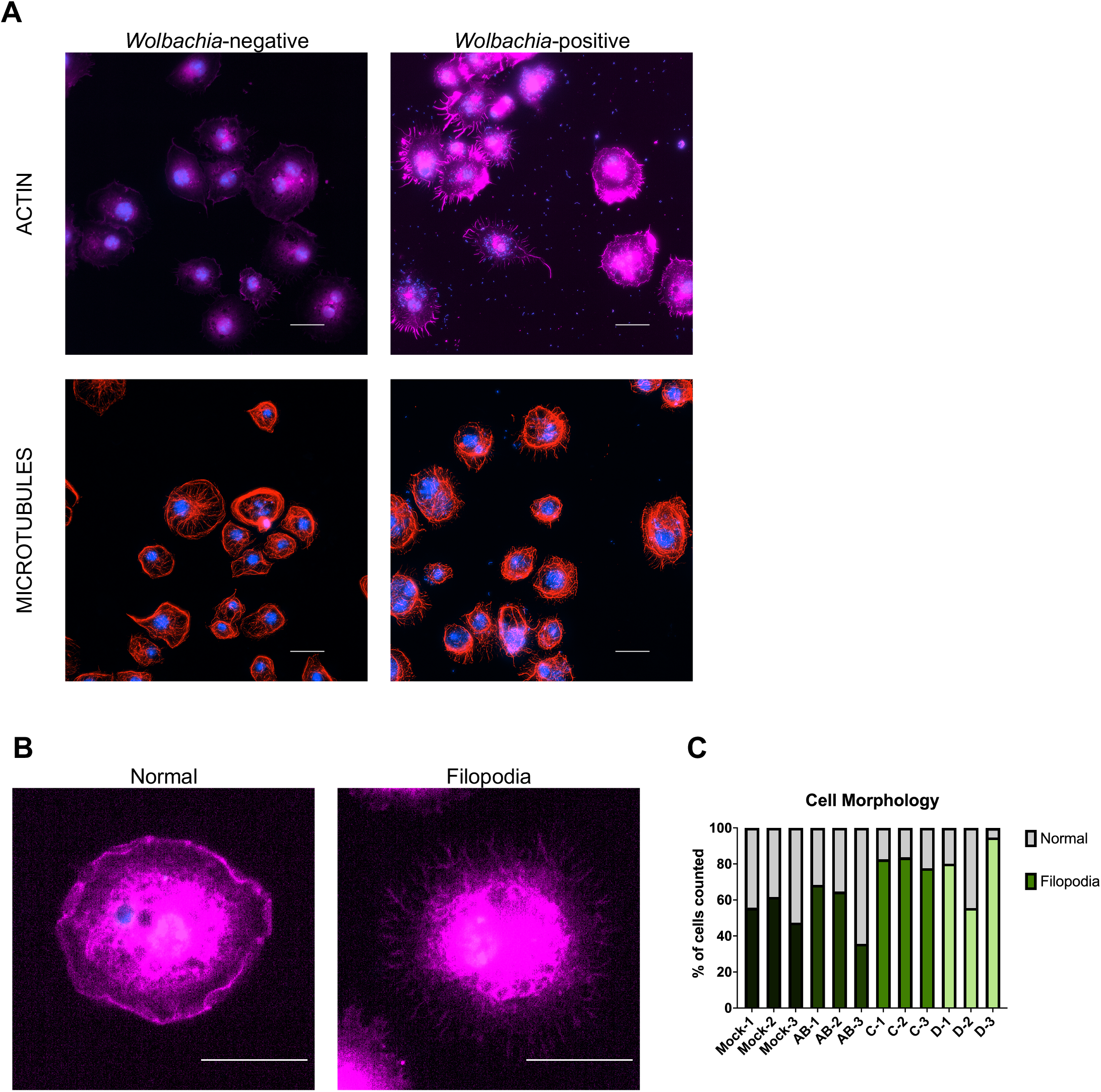
Host cytoskeletal morphology is modified by both *rhoGAP18B* and *Wolbachia*. (A) Host cytoskeletal morphology in the presence and absence of *Wolbachia*. The actin and microtubule networks exhibit different morphology when *Wolbachia* is present in the cell. Since perturbation of these cytoskeletal networks is known to affect the success of *Wolbachia* colonization, it is likely that the endosymbiont manipulates their dynamics to promote its infection. Pink: actin; red: microtubules; blue: DAPI; scale bar: 20 µm. (B) RNAi-mediated knockdown of *rhoGAP18B* isoforms in the absence of *Wolbachia* causes the cell to form filopodia-like structures. Pink: actin; scale bar: 20 µm. (C) Percentage of cells that adopt the filopodia-like morphology when subjected to RNAi-mediated knockdown of *rhoGAP18B*. Each bar is a single replicate. Replicates are defined as populations of cells independently subjected to dsRNA for gene knockdown.

## DISCUSSION

### Wolbachia-mediated viral blocking is robust to host genetic variation in *Drosophila melanogaster*

*Wolbachia*-mediated viral blocking is currently being used to control the spread of arboviral disease in countries where these viruses are endemic. However, climate change and increased temperature regimes might impact *Wolbachia* titer and viral load and transmission, with more variation introduced by host genetic background^13,14,61,62^. Here, we used a set of sequenced *Drosophila* flies to determine if variation in host genetic background influences viral blocking. Uniformly, across the DGRP, the strongest impact of *Wolbachia* colonization was to limit viral replication regardless of fly genotype, indicating that viral blocking is relatively stable and consistent across backgrounds. In contrast, two other studies have previously concluded that variation in host genetic background can impact viral blocking strength. A study from 2019^13^ attained high- and low-blocking host populations through an artificial selection process, which differs from our work here—the DGRP is a collection of flies that were isogenized without this phenotype in mind, and therefore it is possible that this is why we did not observe an impact across the collection. Meanwhile, the study from 2022 demonstrated that a *Wolbachia*-transinfected *Aedes aegypti* mosquito line of Singaporean origin exhibited greater viral blocking strength when compared to one of Mexican origin^14^. In this case, the comparison between only two genetic backgrounds is distinct from our work, which used the power of genetic variation across the entire DGRP to identify variants associated with viral blocking. Although we could certainly cherry-pick a handful of DGRP lines where the range of virus replication with or without *Wolbachia* overlaps, our results did not identify significant genetic variation associated with viral blocking across the dataset in its entirety. Importantly, our results should not be interpreted to mean that host cellular biology does not influence viral blocking, as this has been conclusively explored in the context of null mutations and knockdowns^63–65^. Instead, our results suggest that *Wolbachia-*mediated viral blocking is relatively robust to natural genetic variation, at least in *Drosophila melanogaster*.

### The top GWAS hits for SINV replication are linked through common GTPases

Since host genetic variation led to stark differences in viral replication in the absence of *Wolbachia*, this set was used in a GWAS to reveal specific alleles in genomic regions that contribute to SINV replication. The top hits of this SINV GWAS include *rhoGAP18B* and *betaCOP* (**Figure 3**a). *rhoGAP18B* is a regulator of the actin cytoskeleton, and here we show that loss of isoforms C and CD (but not isoform AB) increased viral replication in the absence of *Wolbachia* (**Figure 4**b), indicating the gene’s anti-viral properties. Meanwhile, *betaCOP* is a component of the COPI coatomer complex, which mediates retrograde Golgi transport^25^, and previous studies have demonstrated that knockdown of this gene results in decreased SINV^66^ and CHIKV^26^, indicating its pro-viral properties.

Interestingly, these two top hits from the GWAS are functionally linked through common GTPases: *rhoGAP18B* is a GTPase activating protein (GAP), and functions to inactivate Rho-family GTPases by promoting their inherent GTP hydrolysis activity. The different isoforms of *rhoGAP18B* act upon distinct Rho-family GTPases: isoform AB binds to Cdc42, isoform D binds to Rac1, and isoform C binds to Cdc42, Rac1, and Rho1^23^. On the other hand, the COPI complex—of which *betaCOP* is a component—recruits activated Cdc42 and Rac (but not Rho), and through the Arp2/3 complex can subsequently polymerize actin at the Golgi, to which the actin binding protein mAbp1 binds and controls trafficking from the ER to the Golgi^28^.

Taken together, the involvement of *rhoGAP18B* and *betaCOP* in SINV replication may be through converging pathways: the anti-viral RhoGAP18B functions to inactivate Rac1, and consequently, inactive Rac1 cannot be recruited to the Golgi by the pro-viral COPI complex. Possibly, the pro-viral nature of *betaCOP* may lie in its promotion of ER-to-Golgi transport: during viral maturation, alphavirus structural proteins are threaded into the ER membrane upon translation, then undergo posttranslational modification as they traverse the secretory pathway from the ER, through the Golgi, to the plasma membrane, from which viral budding occurs^67^. Notably, across the trinity of Rho-family GTPases—Rac1, Rho1, and Cdc42—it seems Rac1 solely connects the cascade in the context of SINV replication since the effect of *rhoGAP18B* knockdown on virus indicates that inactivation of Rac1 and/or Rho1 restricts viral replication, while Cdc42 exerts no effect (**Figure 4**b). Meanwhile, the COPI complex recruits activated Cdc42 and Rac, but not Rho^28^, further narrowing down the list to a single GTPase. Thus, we hypothesize that SINV replication is regulated by a RhoGAP18B-Rac1-betaCOP cascade.

### Actin cytoskeletal organization is the most highly enriched gene set in the SINV GWAS

A gene set enrichment analysis of the loci highlighted by the SINV GWAS indicated that actin cytoskeletal organization was the biological process most strongly associated with viral replication in this dataset. Alphaviruses have been shown to use and manipulate actin for their own benefit, as treatment with cytoskeletal inhibitors result in less internalization of Semliki Forest Virus (inhibition with blebbistatin)^53^, and an increase in Pixuna virus yield (inhibition with cytochalasin D)^60^. Additionally, cells infected with alphavirus form intercellular bridges filled with actin and tubulin^68^; immunostaining revealed that the localization of Semliki Forest Virus nsP3 mimics the striation of filamentous actin^53^; while nsP1 has been shown to depolymerize actin^69^. Although there is some contention in the literature with regards to SINV’s reliance on the actin cytoskeleton^70^, methodological differences in studies could explain these conflicting results. The importance of actin to SINV replication is further bolstered by our results: both *rhoGAP18B* and *betaCOP*—the top hits in the SINV GWAS—are intimately linked with actin, since *rhoGAP18B* is a regulator of actin dynamics, and perturbation of this gene causes morphological changes in the actin cytoskeleton^23^; while the ability of *betaCOP* to mediate ER/Golgi transport is facilitated by its recruitment of Arp2/3, which subsequently leads to actin polymerization^28^.

Actin is also a crossroads where the needs of SINV and *Wolbachia* intersect, as *Wolbachia* has been shown to use actin to achieve successful colonization: flies heterozygous for mutations in the profilin and villin homologs result in decreased *Wolbachia* density^49^. Furthermore, *Wolbachia* manipulates actin, as the *Wolbachia* effector WalE1 is an actin bundler^50^. Thus, because the actin cytoskeleton is intimately involved with both virus and *Wolbachia*, it is possible that actin dynamics during co-infection contribute to viral blocking, as well.

### Prior literature suggests a link between *Wolbachia*-mediated viral blocking and the putative RhoGAP18B-Rac1-betaCOP-actin cascade

Because both *Wolbachia* infection and SINV replication are linked to actin dynamics, and because our GWAS implicated *rhoGAP18B* and *betaCOP* in viral replication, it is possible that the RhoGAP18B-Rac1-betaCOP-actin cascade may also be involved in *Wolbachia*-mediated viral blocking. In support of this, *betaCOP* (as well as several other components of the COPI complex) was found to be a high confidence suppressor of *Wolbachia*: knockdown of the gene resulted in increased *Wolbachia* levels^56^.

However, should this putative cascade be involved in *Wolbachia*-mediated viral blocking, regulation would be through a mechanism distinct from expression modulation, as evidenced by the absence of a transcriptional response from any of the *rhoGAP18B* isoforms in response to the presence of *Wolbachia* (**Supplementary Figure 2**), consistent with our previous work, which also showed no expression differences in both *rhoGAP18B* and *betaCOP* in response to the presence of SINV, *Wolbachia*, or both^59^. However, since key players in this cascade function as molecular switches, they may instead be regulated through in/activation, rather than expression. This is striking, as majority of the most significant SNPs in the *rhoGAP18B* gene region are located in introns (**Table 2**), with many of them falling on transcription factor binding sites previously found to be associated with Dorsal (Dl) and Trithorax-like (Trl)^71^. However, since *rhoGAP18B* is a GAP—a molecular switch—it is more likely that modulation of this gene (whether by the virus or the symbiont) would be through in/activation. Interestingly, the three exonic SNPs of *rhoGAP18B* fall within the C isoform—the isoform for which loss thereof results in an increase in viral replication (**Figure 4**b). These include an insertion (X_19047241; G>GTTG), a deletion (X_19047308; TTGT>T), and a missense mutation (X_19047674; G>A; Ser>Leu). Across these three SNPs, the two fly lines with the lowest viral loads have the major allele (exception: line 1 has no call for the TTGT>T deletion mutation), while the two with the highest viral loads have the variant allele (**Table 2**).

**Table 2.**
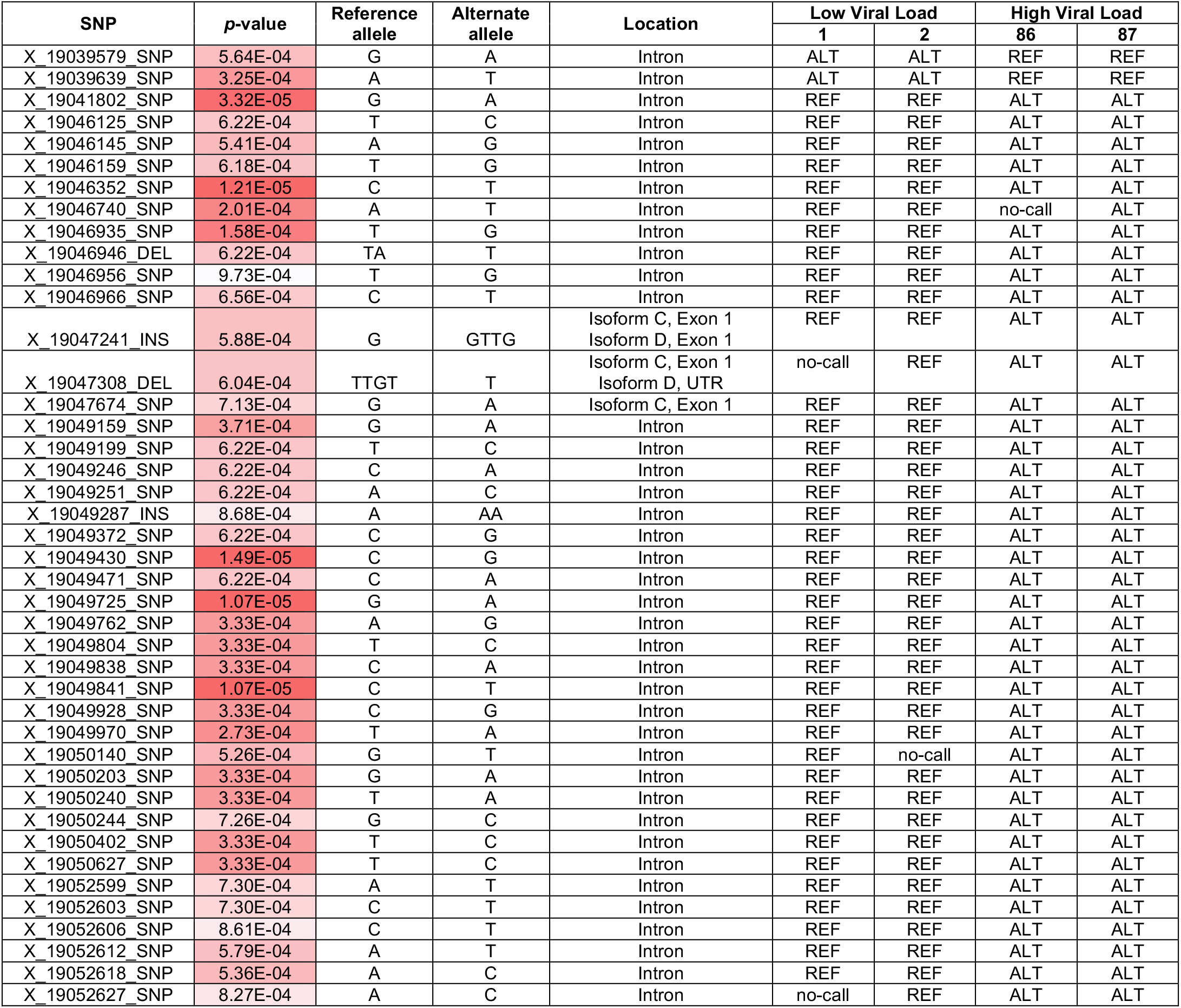
SNPs that fall within the *rhoGAP18B* gene region. All 42 of the highly significant SNPs (*p*<1×10^-3^) within the *rhoGAP18B* gene region, ordered according to their position along the gene. *p*-values are colored according to significance; darker hues represent SNPs of greater significance. For each SNP, the allelic variant found in the fly lines with the lowest and highest viral loads is indicated.

Furthermore, our results show knockdown of *rhoGAP18B* in a *Wolbachia*-positive context did not seem to change the resulting viral load (**Figure 4**b). This could perhaps be due to a myriad of reasons—for one, *Wolbachia* induces such widely-encompassing changes in the cells of its host^13,56,59,72^ that there are likely to be redundancies in the modified pathways that may compensate for the reduction in any one protein. Furthermore, a past study showed that knockdown of *rhoGAP18B* results in morphological changes in the actin cytoskeleton, while overexpression had no visible changes^23^, and as *rhoGAP18B* is a regulator of the actin cytoskeleton, these morphological changes are indicative of function. Interpreting this behavior, it is possible that its cellular function may not need such a large pool of *rhoGAP18B* proteins. With this, it is possible that *Wolbachia* modulates *rhoGAP18B* activity such that even when its amount is diminished, its cellular function to switch off the *betaCOP* cascade is still accomplished, resulting in unperturbed viral restriction.

Future work will determine whether and how SINV and/or *Wolbachia* controls the activation of *rhoGAP18B* and *betaCOP*. For one, RhoGAP18B in cell lines may be replaced with a constitutively active variant using CRISPR, then pull-down assays will measure if the amount of Rac1 recruited by betaCOP is diminished, thus verifying whether the function of these two GWAS hits are indeed linked. For another, the top SNP variants in RhoGAP18B (**Table 2**) can be recapitulated in cell lines, then GTPase pull-down assays will measure how this affects Rac1 activation, thus testing whether SINV modifies RhoGAP18B function.

### Signaling pathways enriched in the GWAS have previously been implicated in viral replication, *Wolbachia* colonization, or *Wolbachia*-mediated viral blocking

The gene set enrichment analysis also revealed that several signaling pathways canonically important in development, cell differentiation, and proliferation are involved in SINV replication (**Supplemental Figure 1** and **Table 3**), and this is supported by a vast array of literature associating these pathways and their components with arboviral replication or the host antiviral immune response. The <u>Notch pathway</u> is linked to host susceptibility to DENV^33^, and a transcriptomic study^34^ has shown the upregulation of several Notch pathway components upon SINV infection—interestingly, two of these upregulated genes (*numb* and *lqf*) are hits in the DGRP GWAS (**Table 3**). The <u>Hippo pathway</u> has been shown to regulate *Drosophila* immunity through the Toll pathway^73^, which is part of the insect antiviral immune response^75^; and *src64B*, a regulator of both the Hippo^92^ and FGF^74^ pathways, is a member of the Src-family kinases, which have been shown to restrict viral replication^76,77^, and have been associated with *Drosophila* immunity^78^. The <u>FGFR pathway</u> is involved in SINV midgut escape in *Aedes aegypti*^35^, and in the suppression of host interferon responses by ZIKV in human fetal astrocytes^36^. While insects lack interferon-based immunity, FGF signaling has been implicated in the *Drosophila* immune response against parasitoid wasp infection^79^—this response occurs through *bnl* signaling (FGF ligand). Although the <u>EGFR pathway</u> has not been directly associated with virus in past literature, it has been linked to FGFR signaling, as these two pathways share *sty* as a common regulator: *sty* regulates FGFR through *bnl*^80,81^, and EGFR through *cbl*^82–84^. Additionally, *cbl* regulates Notch signaling, as well^85,86^. Thus, these pathways are linked to each other through regulators that are all hits in the DGRP GWAS (**Supplemental Figure 1** and **Table 3**).

**Table 3.** Selected gene nodes from the network map of the gene set enrichment analysis. Several genes of interest from the network map, and the number of highly significant SNPs (*p*<1×10^-3^) within their gene region. *p*-value_max_ refers to the *p*-value of the SNP within the indicated gene region that was most significant (top point in the Manhattan plot peak).

| Gene | # of SNPs | $p\text{-value}_{\max}$ | Annotated Pathways |
| --- | --- | --- | --- |
| *numb* | 2 | $2.22 \times 10^{-4}$ | Notch |
| *lqf* | 2 | $6.90 \times 10^{-4}$ | Notch |
| *mnr* | 20 | $6.34 \times 10^{-5}$ | Notch |
| *bnl* | 3 | $9.54 \times 10^{-5}$ | FGFR |
| *sty* | 1 | $4.94 \times 10^{-4}$ | EGFR, FGFR |
| *cbl* | 1 | $9.21 \times 10^{-4}$ | Notch, EGFR |
| *ft* | 6 | $5.87 \times 10^{-5}$ | Hippo |
| *pyd* | 5 | $2.93 \times 10^{-7}$ | Hippo |
| *src64B* | 8 | $5.03 \times 10^{-5}$ | EGFR, Hippo, FGFR |

Interestingly, these pathways are also involved in actin dynamics. The <u>Notch pathway</u> has been shown to activate Rho, leading to phosphorylation of myosin light chain and subsequent reorganization of actin filaments^90^; and activation of PI3/Akt through non-canonical Notch signaling remodels actin through Cdc42^38^. The <u>Hippo pathway</u>^44,45^ both regulates, and is regulated by, actin dynamics^87,88^; *pyd* (which contains the SNP of greatest significance value in the DGRP GWAS), an upstream regulator of Hippo signaling^89^, is a modulator of actin through its binding of cortactin^91^; and the Src-family kinase *src64B* inactivates the Hippo pathway by inducing the accumulation of F-actin^92^ (**Supplemental Figure 1** and **Table 3**). The <u>FGFR pathway</u> upregulates Singed, the *Drosophila* homolog of the actin bundler fascin, allowing for control of actin dynamics during tracheal morphogenesis^43^. Finally, the EGF receptor was reported in 1992 to bind actin^39^, and although more recent work has challenged this finding^41^, the role of the <u>EGFR pathway</u> in actin dynamics has been well-documented^40,42^.

Additionally, some of these pathways are also involved in *Wolbachia* colonization and *Wolbachia*-mediated viral blocking. An artificial selection study in *Aedes aegypti* has associated the <u>Notch pathway</u> with *Wolbachia* blocking of DENV^13,54^, and transcriptomic studies in *Drosophila melanogaster* and *Aedes aegypti* have reported the differential expression of Notch pathway components upon *Wolbachia* colonization^93,94^. Meanwhile, *ft*, an upstream regulator of the <u>Hippo pathway</u>^95,96^, is differentially regulated depending on which strain of *Wolbachia* is colonizing the host^97^ (**Supplemental Figure 1** and **Table 3**).

### Conclusion

In sum, this study suggests that the naturally-occurring variation in host genetic background found in the DGRP does not significantly impact the strength of *Wolbachia*-mediated viral blocking, but does influence SINV replication. The GWAS for SINV replication indicates the involvement of a cascade including RhoGAP18B, Rac1, and betaCOP, leading to the regulation of actin polymerization at the Golgi, which is associated with Golgi trafficking. The similarities between the biological processes highlighted by the SINV GWAS and the cellular impact of *Wolbachia* colonization may indicate a role for the aforementioned cascade in *Wolbachia*-mediated viral blocking. Future work to verify this cascade, along with the specific way by which it abrogates viral replication, will deepen the understanding on the interactions between SINV and the host factors required for its propagation, as well as shed light on the molecular mechanism of *Wolbachia*-mediated viral blocking, which is critical as this endosymbiont continues to be used in practical applications.

## STAR METHODS

### EXPERIMENTAL MODEL AND STUDY PARTICIPANT DETAILS

#### Fly handling

The *Drosophila melanogaster* Reference Panel (DGRP) fly lines used in this study were obtained from the Bloomington *Drosophila* Stock Center. Of the 205 lines, 103 are annotated to be *Wolbachia*-positive^15,16^, but confirmation via qPCR revealed that only 98 had *Wolbachia*, and these were selected for use in this study. These lines were then treated with tetracycline to generate a *Wolbachia*-negative set. All flies were maintained on a 12-hour light/dark cycle at 22°C on BDSC Standard Cornmeal Medium. Mated females used in the screen were age-matched (3 days old) and injected with virus, then subjected to a luciferase assay. Of the 98 DGRP fly lines confirmed to be *Wolbachia*-positive, only 87 were amenable to viral injection.

#### Cell culture

S2 cells were obtained from the *Drosophila* Genomics Resource Center, and JW18 cells were shared by the Sullivan Lab (UC-Santa Cruz). Both cell lines were maintained at 24°C on Schneider’s Insect Medium (Sigma) with 10% heat-inactivated fetal bovine serum (Gibco) and 1% penicillin-streptomycin solution (Gibco). Since JW18 cells are naturally colonized with *Wolbachia*, a *Wolbachia*-negative counterpart was obtained by first treating the cells with a “clearing” dose of tetracycline (10 µg/mL). Once the absence of *Wolbachia* was confirmed, the cells were then cultured on media with a “maintenance” dose of tetracycline (3.3 µg/mL). Prior to experiments, the cells were cultured on media that does not contain tetracycline. BHK-21 cells were obtained from the American Type Culture Collection and maintained at 37°C with 5% CO_2_, on Minimum Essential Eagle’s Medium (Corning) with 10% heat inactivated fetal bovine serum (Corning), 1% antibiotic-antimycotic solution (Corning), 1% L-glutamine (Corning), and 1% non-essential amino acids (Corning).

#### Virus production, injections, and luciferase assays

The virus used in this study is Sindbis virus with nanoluciferase translationally fused to the nsp3 gene (SINV-nsP3-nLuc). Viral RNA was transcribed *in vitro* and transfected into the BHK-21 cells using Lipofectamine LTX (Invitrogen). The virus was propagated, then purified through ultracentrifugation with a 27% sucrose cushion in HNE buffer (20 mM HEPES, 0.15 M NaCl, 0.1 mM EDTA pH 7.5), and resuspended in phosphate-buffered saline.

### METHOD DETAILS

#### Viral injections and luciferase assays

Three-day-old flies were intrathoracically injected with 50 nL of the virus suspension, snap-frozen in liquid nitrogen after three days, and stored at -80°C. Lysis of fly tissues was accomplished using the Luciferase Cell Culture Lysis Reagent (Promega). Luciferase assays were performed using the Nano-Glo Luciferase Assay System (Promega) and read on the BioTek Synergy LX plate reader (for Figure 1) or the Tecan Spark plate reader (for Figure 4).

#### qPCR

To measure *Wolbachia* titer, six-day-old mated female flies were snap-frozen in liquid nitrogen and stored at -80°C until genomic DNA was extracted using the DNeasy Blood & Tissue Kit (Qiagen). qPCR was performed using the iTaq Universal SYBR Green Supermix (Bio-Rad) on the CFX Opus 96 (Bio-Rad). To measure *rhoGAP18B* expression, six-day-old mated female flies were snap-frozen in liquid nitrogen and stored at -80°C until RNA was extracted using the RNeasy Mini Kit (Qiagen). qRT-PCR was performed using the iTaq Universal SYBR Green One-Step Kit (Bio-Rad) on the CFX Opus 96 (Bio-Rad).

#### RNAi-mediated knockdown in cell culture

dsRNA synthesis was performed using the method prescribed by the *Drosophila* RNAi Screening Center^98– 101^. Briefly, RNA was extracted from S2 cells and converted to cDNA using the M-MuLV Reverse Transcriptase (NEB). From this, dsDNA was amplified using the appropriate primer pair with the T7 promoter sequence appended at the 5’ end (5’-TAATACGACTCACTATAGGG), and the Phusion High-Fidelity DNA Polymerase (NEB). The MEGAscript T7 Transcription Kit (ThermoFisher) was then used to transcribe the dsRNA. Finally, the product was purified by RNeasy RNA Purification (Qiagen) and quantified using the Qubit RNA BR Assay Kit (Invitrogen). Cells suspended in serum-free media were treated with 2 µg of dsRNA, and 30 minutes later, supplemented with complete media. After three days, the cells were transferred to coverslips coated with Concanavalin A, allowed to adhere overnight, and subsequently fixed and stained.

#### Fluorescence microscopy

To visualize actin, cells were fixed with 10% formaldehyde in HL3 buffer for 10 minutes at RT, then stained overnight at 4°C with iFluor 594 (1:100; Cayman Chemical). To visualize microtubules, cells were fixed with cold methanol for 5 minutes at -20°C, then stained for 1 hour at RT with the primary antibody DM1α (1:500; Sigma) and for 30 minutes at RT with the secondary antibody Alexa Fluor 594 donkey anti-mouse IgG (H+L) (1:300; Invitrogen). For both protocols, cells were permeabilized and washed using TBS-TX (1% Triton X-100 solution (Sigma) in Tris Buffered Saline); blocked using a solution of 5% bovine serum albumin (Sigma) and 0.1% sodium azide in TBS-TX; stained with 1 µg/mL DAPI for 30 minutes at RT; and mounted in ProLong Diamond (Invitrogen). The coverslips were allowed to cure in the mountant overnight before imaging on the Nikon Ti2 microscope.

### QUANTIFICATION AND STATISTICAL ANALYSIS

Statistical tests (two-way ANOVA with Tukey’s test) and curve-fitting (simple linear regression) were performed on the Prism software by GraphPad (version 10.6.1). Sample sizes (*n* values) are indicated in the corresponding figure legend. Data sets were tested for outliers by the online Outlier Calculator tool by GraphPad with α=0.05 (https://www.graphpad.com/quickcalcs/grubbs1/), and outliers were omitted from plots and analyses. Significance values are represented on plots as asterisks: * = *p*<0.05, ** = *p*<0.01, *** = *p*<0.001, **** = *p*<0.0001. For the GWAS (Figure 3), *p*-values for each SNP were calculated by the DGRPool software.

### KEY RESOURCES TABLE

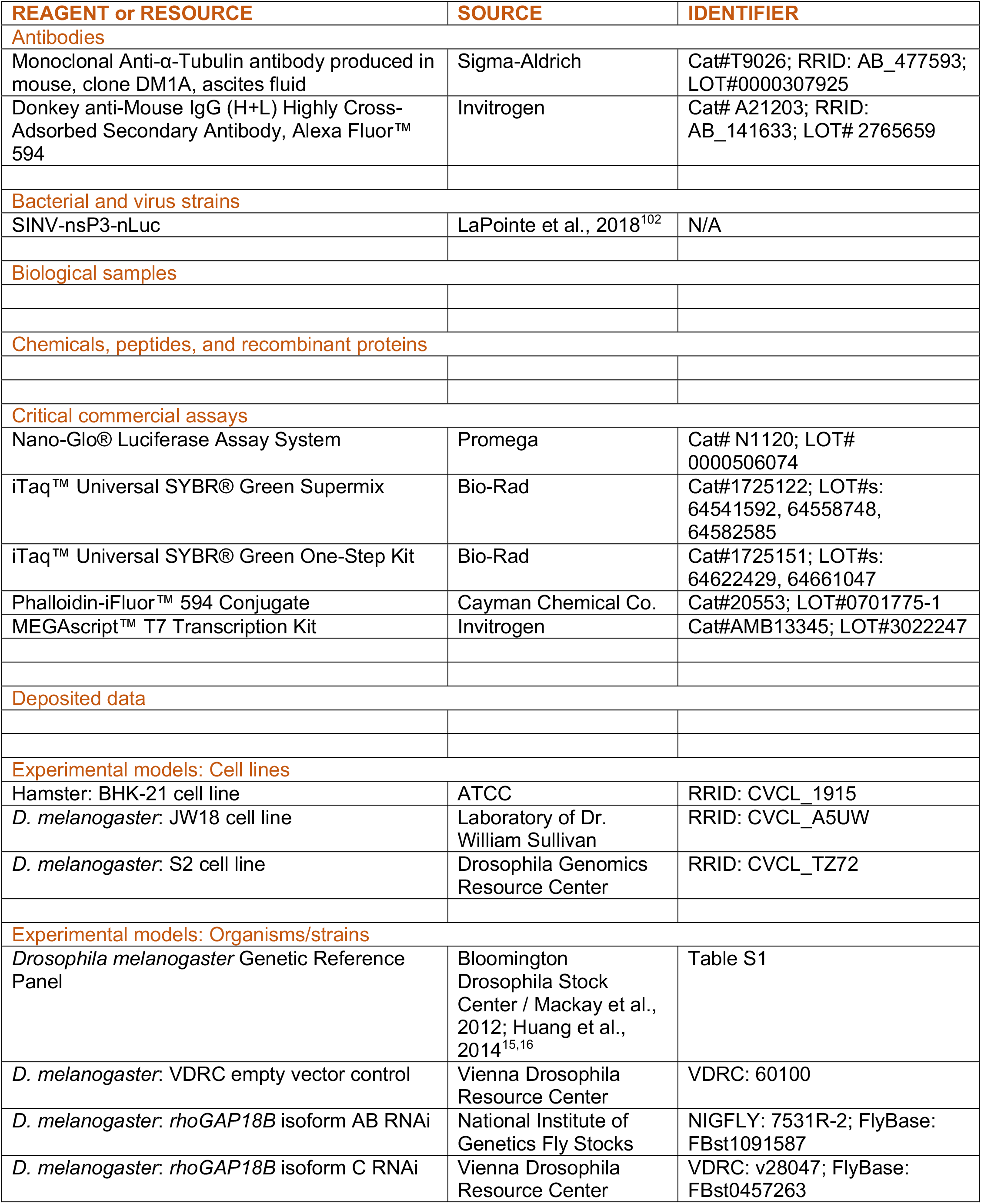
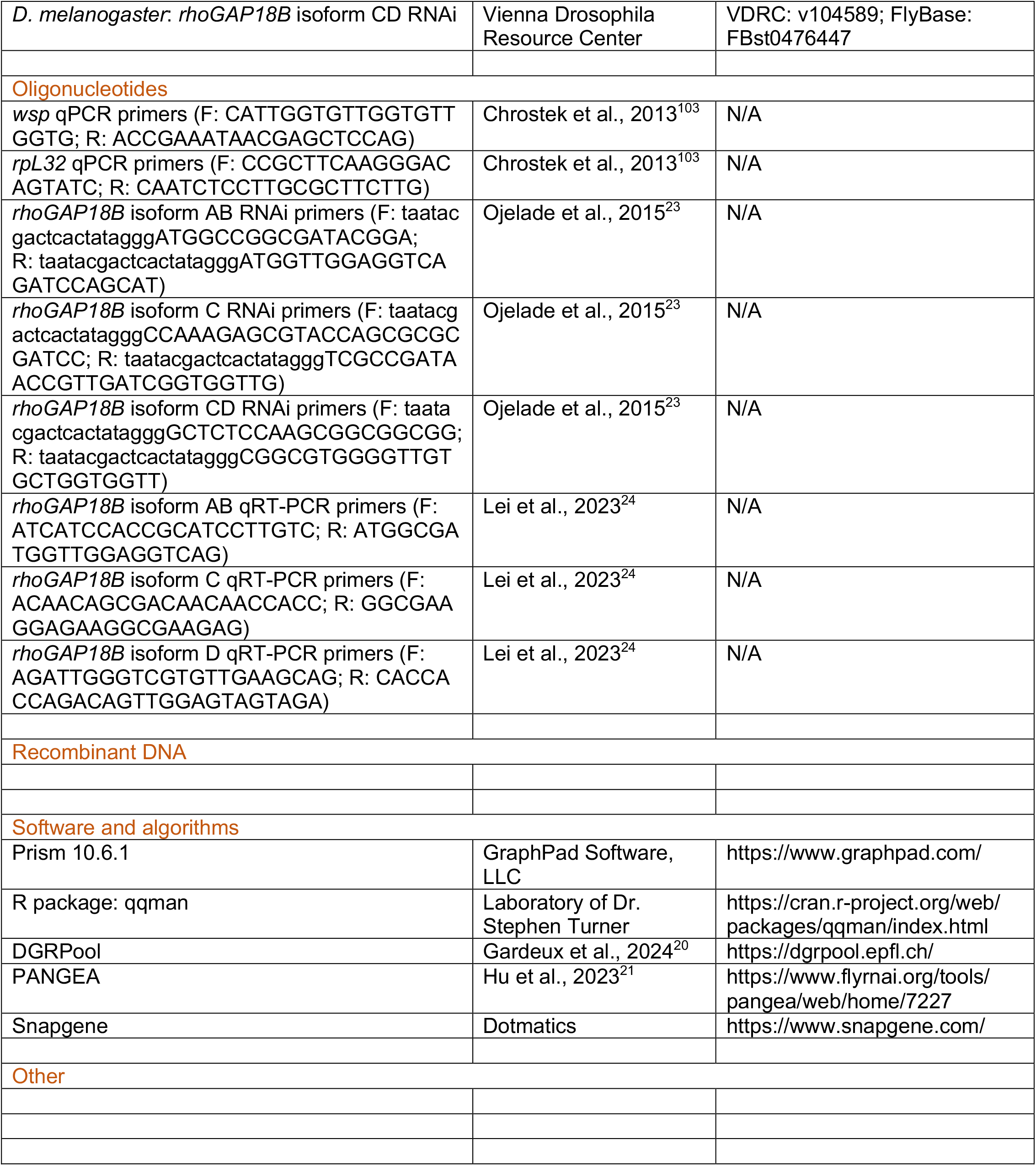

---

**Supplementary Figure 1.**
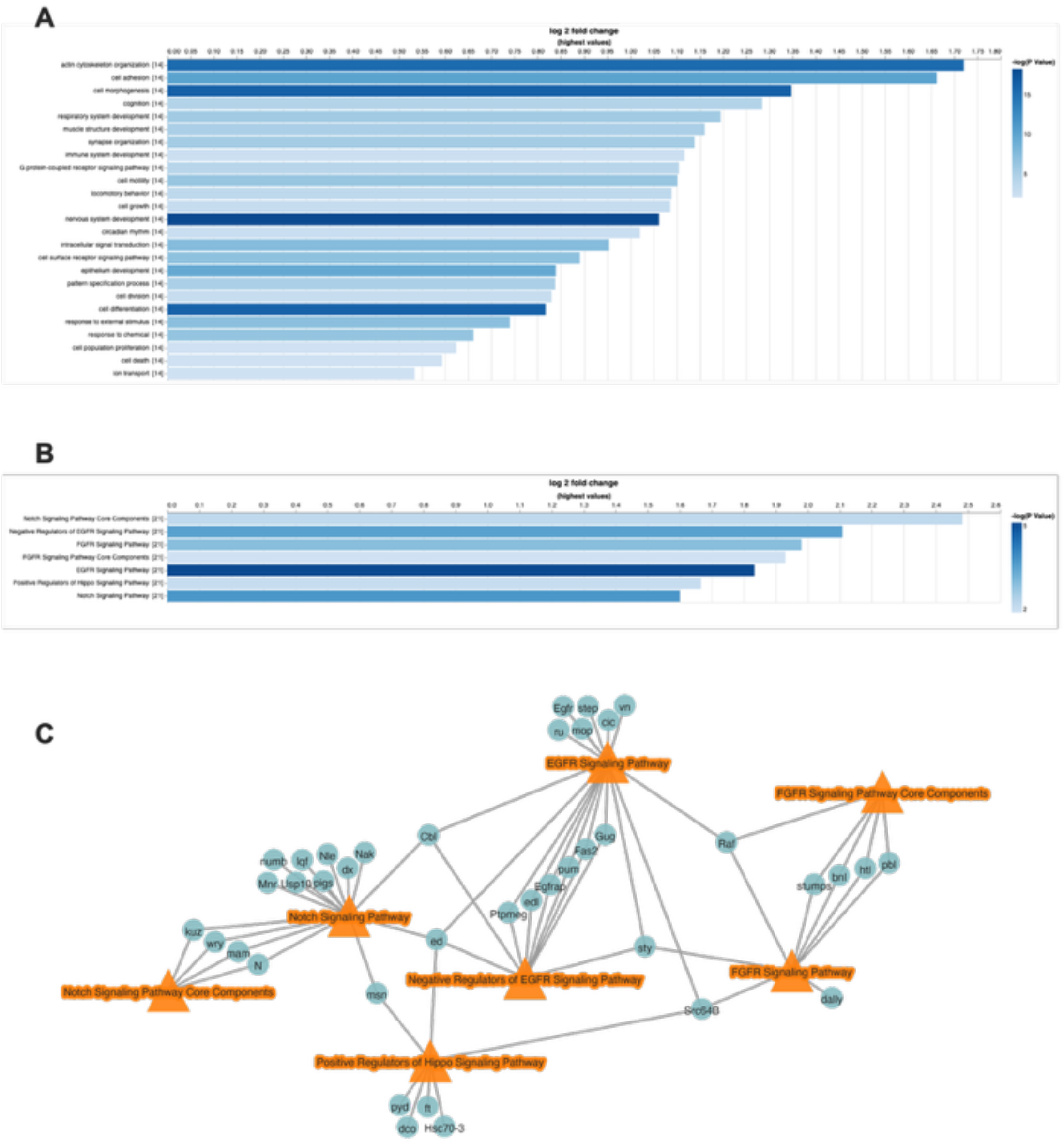
Gene set enrichment analysis of the most significant SNPs from the GWAS. Enriched (A) biological processes and (B) signaling pathways, plotted according to fold change. Darker hues indicate a more significant *p*-value. (C) Enriched signaling pathways visualized on a gene network map, displaying the genes that have been annotated to cross multiple pathways. All gene nodes are hits from the GWAS.

**Supplementary Figure 2.**
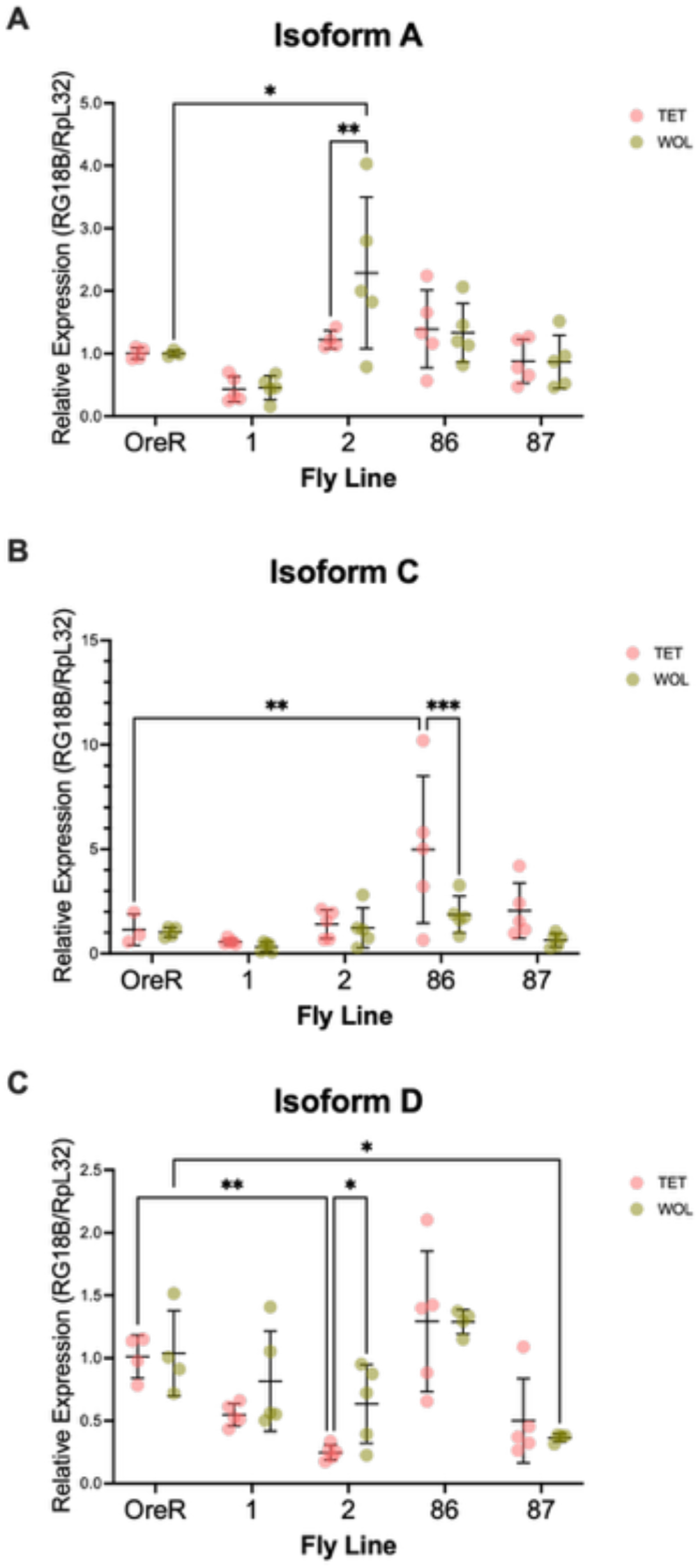
Expression of *rhoGAP18B* in the weakest and strongest blockers, in the presence and absence of *Wolbachia*. Using RT-qPCR, *rhoGAP18B* expression is measured using isoform-specific primers, and in reference to a host housekeeping gene (*rpL32*). Fly lines 1 and 2 are the weakest blockers, fly lines 86 and 87 are the strongest blockers. *Wolbachia*-negative data points in pink, *Wolbachia*-positive data points in green. Each data point corresponds to the expression in a single fly (*n*=5 for each condition).

**Table S1.** Bloomington Drosophila Stock Center identifiers for DGRP lines used.

| **Bloomington Drosophila Stock Center identifier** |
| --- |
| 25177\$ |
| 25183\$ |
| 25186\$ |
| 25187\$ |
| 25190\$ |
| 25195\$ |
| 25198\$ |
| 25200\$ |
| 25201\$ |
| 25202\$ |
| 25208\$ |
| 25209\$ |
| 25210\$ |
| 25445\$ |
| 28122\$ |
| 28130\$ |
| 28131\$ |
| 28142\$ |
| 28144\$ |
| 28145\$ |
| 28146\$ |
| 28149\$ |
| 28151\$ |
| 28152\$ |
| 28156\$ |
| 28164\$ |
| 28165\$ |
| 28168\$ |
| 28172\$ |
| 28173\$ |
| 28174\$ |
| 28178\$ |
| 28180\$ |
| 28182\$ |
| 28185\$ |
| 28189\$ |
| 28190\$ |
| 28197\$ |
| 28198\$ |
| 28208\$ |
| 28212\$ |
| 28213\$ |
| 28215\$ |
| 28219\$ |
| 28220\$ |
| 28223\$ |
| 28224\$ |
| 28227\$ |
| 28229\$ |
| 28230\$ |
| 28233\$ |
| 28234\$ |
| 28235\$ |
| 28236\$ |
| 28237\$ |
| 28242\$ |
| 28243\$ |
| 28244\$ |
| 28245\$ |
| 28249\$ |
| 28250\$ |
| 28251\$ |
| 28253\$ |
| 28254\$ |
| 28255\$ |
| 28256\$ |
| 28257\$ |
| 28258\$ |
| 28260\$ |
| 28276\$ |
| 29651\$ |
| 29654\$ |
| 29655\$ |
| 29656\$ |
| 29659\$ |
| 29660\$ |
| 37525\$ |
| 55016\$ |
| 55018\$ |
| 55023\$ |
| 55024\$ |
| 55025\$ |
| 55026\$ |
| 55032\$ |
| 55038\$ |
| 83728\$ |
| 83729\$ |

## REFERENCES

1. Piantadosi, A., and Kanjilal, S. (2020). Diagnostic Approach for Arboviral Infections in the United States. J. Clin. Microbiol. 58, e01926–19. 10.1128/JCM.01926-19.

2. Schwartz, O., and Albert, M.L. (2010). Biology and pathogenesis of chikungunya virus. Nat. Rev. Microbiol. 8, 491–500. 10.1038/nrmicro2368.

3. Ogunlade, S.T., Meehan, M.T., Adekunle, A.I., Rojas, D.P., Adegboye, O.A., and McBryde, E.S. (2021). A Review: Aedes-Borne Arboviral Infections, Controls and Wolbachia-Based Strategies. Preprint, https://doi.org/10.3390/vaccines9010032 10.3390/vaccines9010032.

4. Whitehorn, J., and Yacoub, S. (2019). Global warming and arboviral infections. Clinical Medicine 19, 149. 10.7861/clinmedicine.19-2-149.

5. Singh, R.K., Dhama, K., Khandia, R., Munjal, A., Karthik, K., Tiwari, R., Chakraborty, S., Malik, Y.S., and Bueno-Marí, R. (2018). Prevention and Control Strategies to Counter Zika Virus, a Special Focus on Intervention Approaches against Vector Mosquitoes—Current Updates. Front. Microbiol. 9.

6. Franz, A.W.E., Sanchez-Vargas, I., Adelman, Z.N., Blair, C.D., Beaty, B.J., James, A.A., and Olson, K.E. (2006). Engineering RNA interference-based resistance to dengue virus type 2 in genetically modified Aedes aegypti. Proceedings of the National Academy of Sciences 103, 4198–4203. 10.1073/pnas.0600479103.

7. Buchman, A., Gamez, S., Li, M., Antoshechkin, I., Li, H.-H., Wang, H.-W., Chen, C.-H., Klein, M.J., Duchemin, J.-B., Crowe Jr., J.E., et al. (2020). Broad dengue neutralization in mosquitoes expressing an engineered antibody. PLoS Pathog. 16, e1008103. 10.1371/journal.ppat.1008103.

8. Lorenzo-Carballa, M.O., Torres-Cambas, Y., Heaton, K., Hurst, G.D.D., Charlat, S., Sherratt, T.N., Van Gossum, H., Cordero-Rivera, A., and Beatty, C.D. (2019). Widespread Wolbachia infection in an insular radiation of damselflies (Odonata, Coenagrionidae). Sci. Rep. 9, 11933. 10.1038/s41598-019-47954-3.

9. Teixeira, L., Ferreira, Á., and Ashburner, M. (2008). The Bacterial Symbiont Wolbachia Induces Resistance to RNA Viral Infections in Drosophila melanogaster. PLoS Biol. 6, e1000002. 10.1371/journal.pbio.1000002.

10. Moreira, L.A., Iturbe-Ormaetxe, I., Jeffery, J.A., Lu, G., Pyke, A.T., Hedges, L.M., Rocha, B.C., Hall-Mendelin, S., Day, A., Riegler, M., et al. (2009). A Wolbachia Symbiont in Aedes aegypti Limits Infection with Dengue, Chikungunya, and Plasmodium. Cell 139, 1268–1278. 10.1016/j.cell.2009.11.042.

11. Hoffmann, A.A., Montgomery, B.L., Popovici, J., Iturbe-Ormaetxe, I., Johnson, P.H., Muzzi, F., Greenfield, M., Durkan, M., Leong, Y.S., Dong, Y., et al. (2011). Successful establishment of Wolbachia in Aedes populations to suppress dengue transmission. Nature 476, 454–457. 10.1038/nature10356.

12. Utarini, A., Indriani, C., Ahmad, R.A., Tantowijoyo, W., Arguni, E., Ansari, M.R., Supriyati, E., Wardana, D.S., Meitika, Y., Ernesia, I., et al. (2021). Efficacy of Wolbachia-Infected Mosquito Deployments for the Control of Dengue. New England Journal of Medicine 384, 2177–2186. 10.1056/NEJMoa2030243.

13. Ford, S.A., Allen, S.L., Ohm, J.R., Sigle, L.T., Sebastian, A., Albert, I., Chenoweth, S.F., and McGraw, E.A. (2019). Selection on Aedes aegypti alters Wolbachia-mediated dengue virus blocking and fitness. Nat. Microbiol. 4, 1832–1839. 10.1038/s41564-019-0533-3.

14. Liang, X., Tan, C.H., Sun, Q., Zhang, M., Wong, P.S.J., Li, M.I., Mak, K.W., Martín-Park, A., Contreras-Perera, Y., Puerta-Guardo, H., et al. (2022). Wolbachia wAlbB remains stable in Aedes aegypti over 15 years but exhibits genetic background-dependent variation in virus blocking. PNAS Nexus 1, pgac203. 10.1093/pnasnexus/pgac203.

15. Mackay, T.F.C., Richards, S., Stone, E.A., Barbadilla, A., Ayroles, J.F., Zhu, D., Casillas, S., Han, Y., Magwire, M.M., Cridland, J.M., et al. (2012). The Drosophila melanogaster Genetic Reference Panel. Nature 482, 173–178. 10.1038/nature10811.

16. Huang, W., Massouras, A., Inoue, Y., Peiffer, J., Ràmia, M., Tarone, A.M., Turlapati, L., Zichner, T., Zhu, D., Lyman, R.F., et al. (2014). Natural variation in genome architecture among 205 Drosophila melanogaster Genetic Reference Panel lines. Genome Res 24, 1193–1208. 10.1101/gr.171546.113.

17. Wang, J.B., Lu, H.-L., and St. Leger, R.J. (2017). The genetic basis for variation in resistance to infection in the Drosophila melanogaster genetic reference panel. PLoS Pathog. 13, e1006260. 10.1371/journal.ppat.1006260.

18. Schmidt, J.M., Battlay, P., Gledhill-Smith, R.S., Good, R.T., Lumb, C., Fournier-Level, A., and Robin, C. (2017). Insights into DDT Resistance from the Drosophila melanogaster Genetic Reference Panel. Genetics 207, 1181–1193. 10.1534/genetics.117.300310.

19. Lu, P., Bian, G., Pan, X., and Xi, Z. (2012). Wolbachia induces density-dependent inhibition to dengue virus in mosquito cells. PLoS Negl Trop Dis 6, e1754. 10.1371/journal.pntd.0001754.

20. Gardeux, V., Bevers, R.P.J., David, F.P.A., Rosschaert, E., Rochepeau, R., and Deplancke, B. (2024). DGRPool: A web tool leveraging harmonized Drosophila Genetic Reference Panel phenotyping data for the study of complex traits. 10.7554/elife.88981.2.

21. Hu, Y., Comjean, A., Attrill, H., Antonazzo, G., Thurmond, J., Chen, W., Li, F., Chao, T., Mohr, S.E., Brown, N.H., et al. (2023). PANGEA: a new gene set enrichment tool for Drosophila and common research organisms. Nucleic Acids Res. 51, W419–W426. 10.1093/nar/gkad331.

22. Clarke, D.N., and Martin, A.C. (2021). Actin-based force generation and cell adhesion in tissue morphogenesis. Current Biology 31, R667–R680. 10.1016/j.cub.2021.03.031.

23. Ojelade, S.A., Acevedo, S.F., Kalahasti, G., Rodan, A.R., and Rothenfluh, A. (2015). RhoGAP18B Isoforms Act on Distinct Rho-Family GTPases and Regulate Behavioral Responses to Alcohol via Cofilin. PLoS One 10, e0137465. 10.1371/journal.pone.0137465.

24. Lei, F., Xu, X., Huang, J., Su, D., and Wan, P. (2023). Drosophila RhoGAP18B regulates actin cytoskeleton during border cell migration. PLoS One 18, e0280652. 10.1371/journal.pone.0280652.

25. Cottam, N.P., and Ungar, D. (2012). Retrograde vesicle transport in the Golgi. Protoplasma 249, 943–955. 10.1007/s00709-011-0361-7.

26. Zhang, N., and Zhang, L. (2017). Key components of COPI and COPII machineries are required for chikungunya virus replication. Biochem. Biophys. Res. Commun. 493, 1190–1196. 10.1016/j.bbrc.2017.09.142.

27. Wu, W.J., Erickson, J.W., Lin, R., and Cerione, R.A. (2000). The γ-subunit of the coatomer complex binds Cdc42 to mediate transformation. Nature 405, 800–804. 10.1038/35015585.

28. Fucini, R. V, Chen, J.-L., Sharma, C., Kessels, M.M., and Stamnes, M. (2002). Golgi Vesicle Proteins Are Linked to the Assembly of an Actin Complex Defined by mAbp1. Mol. Biol. Cell 13, 621–631. 10.1091/mbc.01-11-0547.

29. Zhou, B., Lin, W., Long, Y., Yang, Y., Zhang, H., Wu, K., and Chu, Q. (2022). Notch signaling pathway: architecture, disease, and therapeutics. Signal Transduct. Target. Ther. 7, 95. 10.1038/s41392-022-00934-y.

30. Xie, Y., Su, N., Yang, J., Tan, Q., Huang, S., Jin, M., Ni, Z., Zhang, B., Zhang, D., Luo, F., et al. (2020). FGF/FGFR signaling in health and disease. Signal Transduct. Target. Ther. 5, 181. 10.1038/s41392-020-00222-7.

31. Bogdan, S., and Klämbt, C. (2001). Epidermal growth factor receptor signaling. Current Biology 11, R292–R295. 10.1016/S0960-9822(01)00167-1.

32. Fu, M., Hu, Y., Lan, T., Guan, K.-L., Luo, T., and Luo, M. (2022). The Hippo signalling pathway and its implications in human health and diseases. Signal Transduct. Target. Ther. 7, 376. 10.1038/s41392-022-01191-9.

33. Taracena, M.L., Bottino-Rojas, V., Talyuli, O.A.C., Walter-Nuno, A.B., Oliveira, J.H.M., Angleró-Rodriguez, Y.I., Wells, M.B., Dimopoulos, G., Oliveira, P.L., and Paiva-Silva, G.O. (2018). Regulation of midgut cell proliferation impacts Aedes aegypti susceptibility to dengue virus. PLoS Negl. Trop. Dis. 12, e0006498. 10.1371/journal.pntd.0006498.

34. Mudiganti, U., Hernandez, R., and Brown, D.T. (2010). Insect response to alphavirus infection— Establishment of alphavirus persistence in insect cells involves inhibition of viral polyprotein cleavage. Virus Res. 150, 73–84. 10.1016/j.virusres.2010.02.016.

35. Wu, W., Simmons, C.A., Moffitt, J., Clem, R.J., and Passarelli, A.L. (2020). Effects of Manipulating Fibroblast Growth Factor Expression on Sindbis Virus Replication In Vitro and in Aedes aegypti Mosquitoes. Viruses 12. 10.3390/v12090943.

36. Limonta, D., Jovel, J., Kumar, A., Lu, J., Hou, S., Airo, A.M., Lopez-Orozco, J., Wong, C.P., Saito, L., Branton, W., et al. (2019). Fibroblast Growth Factor 2 Enhances Zika Virus Infection in Human Fetal Brain. J. Infect. Dis. 220, 1377–1387. 10.1093/infdis/jiz073.

37. Garcia Jr., G., Paul, S., Beshara, S., Ramanujan, V.K., Ramaiah, A., Nielsen-Saines, K., Li, M.M.H., French, S.W., Morizono, K., Kumar, A., et al. (2020). Hippo Signaling Pathway Has a Critical Role in Zika Virus Replication and in the Pathogenesis of Neuroinflammation. Am. J. Pathol. 190, 844–861. 10.1016/j.ajpath.2019.12.005.

38. Liu, L., Zhang, L., Zhao, S., Zhao, X.-Y., Min, P.-X., Ma, Y.-D., Wang, Y.-Y., Chen, Y., Tang, S.-J., Zhang, Y.-J., et al. (2019). Non-canonical Notch Signaling Regulates Actin Remodeling in Cell Migration by Activating PI3K/AKT/Cdc42 Pathway. Front. Pharmacol. Volume 10-2019.

39. den Hartigh, J.C., van Bergen en Henegouwen, P.M., Verkleij, A.J., and Boonstra, J. (1992). The EGF receptor is an actin-binding protein. Journal of Cell Biology 119, 349–355. 10.1083/jcb.119.2.349.

40. Singhai, A., Wakefield, D.L., Bryant, K.L., Hammes, S.R., Holowka, D., and Baird, B. (2014). Spatially Defined EGF Receptor Activation Reveals an F-Actin-Dependent Phospho-Erk Signaling Complex. Biophys. J. 107, 2639–2651. 10.1016/j.bpj.2014.09.048.

41. Pandey, S., and Wohland, T. (2024). EGFR does not directly interact with cortical actin: A&#xa0;SRRF’n’TIRF study. Biophys. J. 123, 3736–3749. 10.1016/j.bpj.2024.09.022.

42. Clarke, D.N., Miller, P.W., and Martin, A.C. (2025). EGFR-dependent actomyosin patterning coordinates morphogenetic movements between tissues in <em>Drosophila melanogaster</em>. Dev. Cell 60, 270–287.e6. 10.1016/j.devcel.2024.10.002.

43. Okenve-Ramos, P., and Llimargas, M. (2014). Fascin links Btl/FGFR signalling to the actin cytoskeleton during Drosophila tracheal morphogenesis. Development 141, 929–939. 10.1242/dev.103218.

44. Matsui, Y., and Lai, Z.-C. (2013). Mutual regulation between Hippo signaling and actin cytoskeleton. Protein Cell 4, 904–910. 10.1007/s13238-013-3084-z.

45. Seo, J., and Kim, J. (2018). Regulation of Hippo signaling by actin remodeling. BMB Rep. 51, 151–156. 10.5483/BMBRep.2018.51.3.012.

46. Bernards, A. (2003). GAPs galore! A survey of putative Ras superfamily GTPase activating proteins in man and Drosophila. Biochimica et Biophysica Acta (BBA) - Reviews on Cancer 1603, 47–82. 10.1016/S0304-419X(02)00082-3.

47. Rothenfluh, A., Threlkeld, R.J., Bainton, R.J., Tsai, L.T.-Y., Lasek, A.W., and Heberlein, U. (2006). Distinct Behavioral Responses to Ethanol Are Regulated by Alternate RhoGAP18B Isoforms. Cell 127, 199–211. 10.1016/j.cell.2006.09.010.

48. Kamath, A.D., Deehan, M.A., and Frydman, H.M. (2018). Polar cell fate stimulates Wolbachia intracellular growth. Development 145, dev158097. 10.1242/dev.158097.

49. Newton, I.L.G., Savytskyy, O., and Sheehan, K.B. (2015). Wolbachia Utilize Host Actin for Efficient Maternal Transmission in Drosophila melanogaster. PLoS Pathog. 11, e1004798. 10.1371/journal.ppat.1004798.

50. Sheehan Kathy, B., Martin, M., Lesser Cammie, F., Isberg Ralph, R., and Newton Irene, L.G. (2016). Identification and Characterization of a Candidate Wolbachia pipientis Type IV Effector That Interacts with the Actin Cytoskeleton. mBio 7, e00622–16. 10.1128/mBio.00622-16.

51. Martin, M., López-Madrigal, S., and Newton, I.L.G. (2024). The Wolbachia WalE1 effector alters Drosophila endocytosis. PLoS Pathog. 20, e1011245..

52. Geoghegan, V., Stainton, K., Rainey, S.M., Ant, T.H., Dowle, A.A., Larson, T., Hester, S., Charles, P.D., Thomas, B., and Sinkins, S.P. (2017). Perturbed cholesterol and vesicular trafficking associated with dengue blocking in Wolbachia-infected Aedes aegypti cells. Nat. Commun. 8, 526. 10.1038/s41467-017-00610-8.

53. Spuul, P., Balistreri, G., Kääriäinen, L., and Ahola, T. (2010). Phosphatidylinositol 3-Kinase-, Actin-, and Microtubule-Dependent Transport of Semliki Forest Virus Replication Complexes from the Plasma Membrane to Modified Lysosomes. J. Virol. 84, 7543–7557. 10.1128/jvi.00477-10.

54. Ford, S.A., Albert, I., Allen, S.L., Chenoweth, S.F., Jones, M., Koh, C., Sebastian, A., Sigle, L.T., and McGraw, E.A. (2020). Artificial Selection Finds New Hypotheses for the Mechanism of Wolbachia-Mediated Dengue Blocking in Mosquitoes. Front. Microbiol. Volume 11-2020.

55. Sigle, L.T., Jones, M., Novelo, M., Ford, S.A., Urakova, N., Lymperopoulos, K., Sayre, R.T., Xi, Z., Rasgon, J.L., and McGraw, E.A. (2022). Assessing Aedes aegypti candidate genes during viral infection and Wolbachia-mediated pathogen blocking. Insect Mol. Biol. 31, 356–368. 10.1111/imb.12764.

56. Grobler, Y., Yun, C.Y., Kahler, D.J., Bergman, C.M., Lee, H., Oliver, B., and Lehmann, R. (2018). Whole genome screen reveals a novel relationship between Wolbachia levels and Drosophila host translation. PLoS Pathog. 14, e1007445..

57. Yun-heng, M., Wei-hao, D., Jing, L., Da-wei, H., and Jin-hua, X. (2024). Single-cell transcriptome sequencing reveals that Wolbachia induces gene expression changes in Drosophila ovary cells to favor its own maternal transmission. mBio 15, e01473–24. 10.1128/mbio.01473-24.

58. Radoshitzky, S.R., Pegoraro, G., Chī, X., Dǒng, L., Chiang, C.-Y., Jozwick, L., Clester, J.C., Cooper, C.L., Courier, D., Langan, D.P., et al. (2016). siRNA Screen Identifies Trafficking Host Factors that Modulate Alphavirus Infection. PLoS Pathog. 12, e1005466. 10.1371/journal.ppat.1005466.

59. Lindsey Amelia, R.I., Bhattacharya, T., Hardy Richard, W., and Newton Irene, L.G. (2021). Wolbachia and Virus Alter the Host Transcriptome at the Interface of Nucleotide Metabolism Pathways. mBio 12, e03472–20. 10.1128/mBio.03472-20.

60. Gil, P.I., Albrieu-Llinás, G., Mlewski, E.C., Monetti, M., Fozzatti, L., Cuffini, C., Fernández Romero, J., Kunda, P., and Paglini, M.G. (2017). Pixuna virus modifies host cell cytoskeleton to secure infection. Sci. Rep. 7, 5757. 10.1038/s41598-017-05983-w.

61. Ross, P.A., Elfekih, S., Collier, S., Klein, M.J., Lee, S.S., Dunn, M., Jackson, S., Zhang, Y., Axford, J.K., Gu, X., et al. (2023). Developing Wolbachia-based disease interventions for an extreme environment. PLoS Pathog. 19, e1011117..

62. Murdock, C.C., Blanford, S., Hughes, G.L., Rasgon, J.L., and Thomas, M.B. (2014). Temperature alters Plasmodium blocking by Wolbachia. Sci. Rep. 4, 3932. 10.1038/srep03932.

63. Bhattacharya, T., Newton, I.L.G., and Hardy, R.W. (2017). Wolbachia elevates host methyltransferase expression to block an RNA virus early during infection. PLoS Pathog. 13, e1006427. 10.1371/journal.ppat.1006427.

64. Urakova, N., Joseph, R.E., Huntsinger, A., Macias, V.M., Jones, M.J., Sigle, L.T., Li, M., Akbari, O.S., Xi, Z., Lymperopoulos, K., et al. (2024). Alpha-mannosidase-2 modulates arbovirus infection in a pathogen- and Wolbachia-specific manner in Aedes aegypti mosquitoes. Insect Mol. Biol. 33, 362–371. 10.1111/imb.12904.

65. Nicole, S., Ram, P., Lauren, G., Adela, K., B, R.D., G, N.I.L., and W, H.R. (2025). Pseudouridine synthases are proviral factors for Sindbis virus in insect and mammalian cells. mBio 16, e01329–25. 10.1128/mbio.01329-25.

66. Panda, D., Rose, P.P., Hanna, S.L., Gold, B., Hopkins, K.C., Lyde, R.B., Marks, M.S., and Cherry, S. (2013). Genome-wide RNAi Screen Identifies SEC61A and VCP as Conserved Regulators of Sindbis Virus Entry. Cell Rep. 5, 1737–1748. 10.1016/j.celrep.2013.11.028.

67. Brown, R.S., Wan, J.J., and Kielian, M. (2018). The Alphavirus Exit Pathway: What We Know and What We Wish We Knew. Viruses 10. 10.3390/v10020089.

68. Martinez, M.G., and Kielian, M. (2016). Intercellular Extensions Are Induced by the Alphavirus Structural Proteins and Mediate Virus Transmission. PLoS Pathog. 12, e1006061..

69. Laakkonen, P., Auvinen, P., Kujala, P., and Kääriäinen, L. (1998). Alphavirus Replicase Protein NSP1 Induces Filopodia and Rearrangement of Actin Filaments. J. Virol. 72, 10265–10269. 10.1128/jvi.72.12.10265-10269.1998.

70. Matthews, J.D., Morgan, R., Sleigher, C., and Frey, T.K. (2013). Do viruses require the cytoskeleton? Virol. J. 10, 121. 10.1186/1743-422X-10-121.

71. Consortium, T. modENCODE, Roy, S., Ernst, J., Kharchenko, P. V, Kheradpour, P., Negre, N., Eaton, M.L., Landolin, J.M., Bristow, C.A., Ma, L., et al. (2010). Identification of Functional Elements and Regulatory Circuits by Drosophila modENCODE. Science (1979). 330, 1787–1797. 10.1126/science.1198374.

72. White, P.M., Serbus, L.R., Debec, A., Codina, A., Bray, W., Guichet, A., Lokey, R.S., and Sullivan, W. (2017). Reliance of Wolbachia on High Rates of Host Proteolysis Revealed by a Genome-Wide RNAi Screen of Drosophila Cells. Genetics 205, 1473–1488. 10.1534/genetics.116.198903.

73. Liu, B., Zheng, Y., Yin, F., Yu, J., Silverman, N., and Pan, D. (2016). Toll Receptor-Mediated Hippo Signaling Controls Innate Immunity in Drosophila. Cell 164, 406–419. 10.1016/j.cell.2015.12.029.

74. Muha, V., and Müller, H.-A.J. (2013). Functions and Mechanisms of Fibroblast Growth Factor (FGF) Signalling in Drosophila melanogaster. Int. J. Mol. Sci. 14, 5920–5937. 10.3390/ijms14035920.

75. Kingsolver, M.B., Huang, Z., and Hardy, R.W. (2013). Insect Antiviral Innate Immunity: Pathways, Effectors, and Connections. J. Mol. Biol. 425, 4921–4936. 10.1016/j.jmb.2013.10.006.

76. Rebecca, B., Sanjay, S. A M.N., Jennifer, T., N, K.C., Patricia, S., Lee, G., R, D.V., T, H.M., E, M.T., et al. (2019). Src Family Kinase Inhibitors Block Translation of Alphavirus Subgenomic mRNAs. Antimicrob. Agents Chemother. 63, 10.1128/aac.02325-18. 10.1128/aac.02325-18.

77. Wei, M., Zhang, Y., Aweya, J.J., Wang, F., Li, S., Lun, J., Zhu, C., and Yao, D. (2019). Litopenaeus vannamei Src64B restricts white spot syndrome virus replication by modulating apoptosis. Fish Shellfish Immunol. 93, 313–321. 10.1016/j.fsi.2019.07.062.

78. Unckless, R.L., Rottschaefer, S.M., and Lazzaro, B.P. (2015). The Complex Contributions of Genetics and Nutrition to Immunity in Drosophila melanogaster. PLoS Genet. 11, e1005030..

79. Tattikota, S.G., Cho, B., Liu, Y., Hu, Y., Barrera, V., Steinbaugh, M.J., Yoon, S.-H., Comjean, A., Li, F., Dervis, F., et al. (2020). A single-cell survey of Drosophila blood. Elife 9, e54818. 10.7554/eLife.54818.

80. Hacohen, N., Kramer, S., Sutherland, D., Hiromi, Y., and Krasnow, M.A. (1998). <em>sprouty</em> Encodes a Novel Antagonist of FGF Signaling that Patterns Apical Branching of the <em>Drosophila</em> Airways. Cell 92, 253–263. 10.1016/S0092-8674(00)80919-8.

81. Jarvis, L.A., Toering, S.J., Simon, M.A., Krasnow, M.A., and Smith-Bolton, R.K. (2006). Sprouty proteins are in vivo targets of Corkscrew/SHP-2 tyrosine phosphatases. Development 133, 1133–1142. 10.1242/dev.02255.

82. Wong, E.S.M., Lim, J., Low, B.C., Chen, Q., and Guy, G.R. (2001). Evidence for Direct Interaction between Sprouty and Cbl *. Journal of Biological Chemistry 276, 5866–5875. 10.1074/jbc.M006945200.

83. Wang, P.-Y., and Pai, L.-M. (2011). D-Cbl Binding to Drk Leads to Dose-Dependent Down-Regulation of EGFR Signaling and Increases Receptor-Ligand Endocytosis. PLoS One 6, e17097..

84. Egan, J.E., Hall, A.B., Yatsula, B.A., and Bar-Sagi, D. (2002). The bimodal regulation of epidermal growth factor signaling by human Sprouty proteins. Proceedings of the National Academy of Sciences 99, 6041–6046. 10.1073/pnas.052090899.

85. Wang, Y., Chen, Z., and Bergmann, A. (2010). Regulation of EGFR and Notch signaling by distinct isoforms of D-cbl during Drosophila development. Dev. Biol. 342, 1–10. 10.1016/j.ydbio.2010.03.005.

86. Bala Tannan, N., Collu, G., Humphries, A.C., Serysheva, E., Weber, U., and Mlodzik, M. (2018). AKAP200 promotes Notch stability by protecting it from Cbl/lysosome-mediated degradation in Drosophila melanogaster. PLoS Genet. 14, e1007153..

87. Sansores‐Garcia, L., Bossuyt, W., Wada, K., Yonemura, S., Tao, C., Sasaki, H., and Halder, G. (2011). Modulating F‐actin organization induces organ growth by affecting the Hippo pathway. EMBO J. 30, 2325–2335–2335. 10.1038/emboj.2011.157.

88. Lucas, E.P., Khanal, I., Gaspar, P., Fletcher, G.C., Polesello, C., Tapon, N., and Thompson, B.J. (2013). The Hippo pathway polarizes the actin cytoskeleton during collective migration of Drosophila border cells. Journal of Cell Biology 201, 875–885. 10.1083/jcb.201210073.

89. Sang, Q., Wang, G., Morton, D.B., Wu, H., and Xie, B. (2021). The ZO-1 protein Polychaetoid as an upstream regulator of the Hippo pathway in Drosophila. PLoS Genet. 17, e1009894..

90. Venkatesh, D., Fredette, N., Rostama, B., Tang, Y., Vary, C.P.H., Liaw, L., and Urs, S. (2011). RhoA-Mediated Signaling in Notch-Induced Senescence-Like Growth Arrest and Endothelial Barrier Dysfunction. Arterioscler. Thromb. Vasc. Biol. 31, 876–882. 10.1161/ATVBAHA.110.221945.

91. Katsube, T., Takahisa, M., Ueda, R., Hashimoto, N., Kobayashi, M., and Togashi, S. (1998). Cortactin Associates with the Cell-Cell Junction Protein ZO-1 in both <em>Drosophila</em> and Mouse *. Journal of Biological Chemistry 273, 29672–29677. 10.1074/jbc.273.45.29672.

92. Enomoto, M., and Igaki, T. (2013). Src controls tumorigenesis via JNK‐dependent regulation of the Hippo pathway in *Drosophila*. EMBO Rep. 14, 65–72–72. 10.1038/embor.2012.185.

93. Lindsey, A.R.I., Tennessen, J.M., Gelaw, M.A., Jones, M.W., Parish, A.J., Newton, I.L.G., Nemkov, T., D’Alessandro, A., Rai, M., and Stark, N. (2025). The intracellular symbiont Wolbachia alters Drosophila development and metabolism to buffer against nutritional stress. PLoS Genet. 21, e1011905..

94. Wimalasiri-Yapa, B.M.C.R., Huang, B., Ross, P.A., Hoffmann, A.A., Ritchie, S.A., Frentiu, F.D., Warrilow, D., and van den Hurk, A.F. (2023). Differences in gene expression in field populations of Wolbachia-infected Aedes aegypti mosquitoes with varying release histories in northern Australia. PLoS Negl. Trop. Dis. 17, e0011222..

95. Bennett, F.C., and Harvey, K.F. (2006). Fat Cadherin Modulates Organ Size in <em>Drosophila</em> via the Salvador/Warts/Hippo Signaling Pathway. Current Biology 16, 2101–2110. 10.1016/j.cub.2006.09.045.

96. Willecke, M., Hamaratoglu, F., Kango-Singh, M., Udan, R., Chen, C., Tao, C., Zhang, X., and Halder, G. (2006). The Fat Cadherin Acts through the Hippo Tumor-Suppressor Pathway to Regulate Tissue Size. Current Biology 16, 2090–2100. 10.1016/j.cub.2006.09.005.

97. Gruntenko, N.E., Deryuzhenko, M.A., Andreenkova, O. V, Shishkina, O.D., Bobrovskikh, M.A., Shatskaya, N. V, and Vasiliev, G. V (2023). Drosophila melanogaster Transcriptome Response to Different Wolbachia Strains. Int. J. Mol. Sci. 24. 10.3390/ijms242417411.

98. Clemens, J.C., Worby, C.A., Simonson-Leff, N., Muda, M., Maehama, T., Hemmings, B.A., and Dixon, J.E. (2000). Use of double-stranded RNA interference in Drosophila cell lines to dissect signal transduction pathways. Proceedings of the National Academy of Sciences 97, 6499–6503. 10.1073/pnas.110149597.

99. Armknecht, S., Boutros, M., Kiger, A., Nybakken, K., Mathey-Prevot, B., and Perrimon, N. (2005). High-Throughput RNA Interference Screens in Drosophila Tissue Culture Cells∗∗S. A., M. B., and A. K. contributed equally to this work. In Methods in Enzymology (Academic Press), pp. 55–73. 10.1016/S0076-6879(04)92004-6.

100. Flockhart, I., Booker, M., Kiger, A., Boutros, M., Armknecht, S., Ramadan, N., Richardson, K., Xu, A., Perrimon, N., and Mathey-Prevot, B. (2006). FlyRNAi: the Drosophila RNAi screening center database. Nucleic Acids Res. 34, D489–D494. 10.1093/nar/gkj114.

101. Flockhart, I.T., Booker, M., Hu, Y., McElvany, B., Gilly, Q., Mathey-Prevot, B., Perrimon, N., and Mohr, S.E. (2012). FlyRNAi.org—the database of the Drosophila RNAi screening center: 2012 update. Nucleic Acids Res. 40, D715–D719. 10.1093/nar/gkr953.

102. T, L.A., Joaquín, M.-C., and J, S.K. (2018). Increasing the Capping Efficiency of the Sindbis Virus nsP1 Protein Negatively Affects Viral Infection. mBio 9, 10.1128/mbio.02342-18. 10.1128/mbio.02342-18.

103. Chrostek, E., Marialva, M.S.P., Esteves, S.S., Weinert, L.A., Martinez, J., Jiggins, F.M., and Teixeira, L. (2013). Wolbachia Variants Induce Differential Protection to Viruses in Drosophila melanogaster: A Phenotypic and Phylogenomic Analysis. PLoS Genet. 9, e1003896..

