## Supplementary figures and images for "Skeletons in the closet: The importance of actin in alphavirus replication"

### Supplementary Figure 1

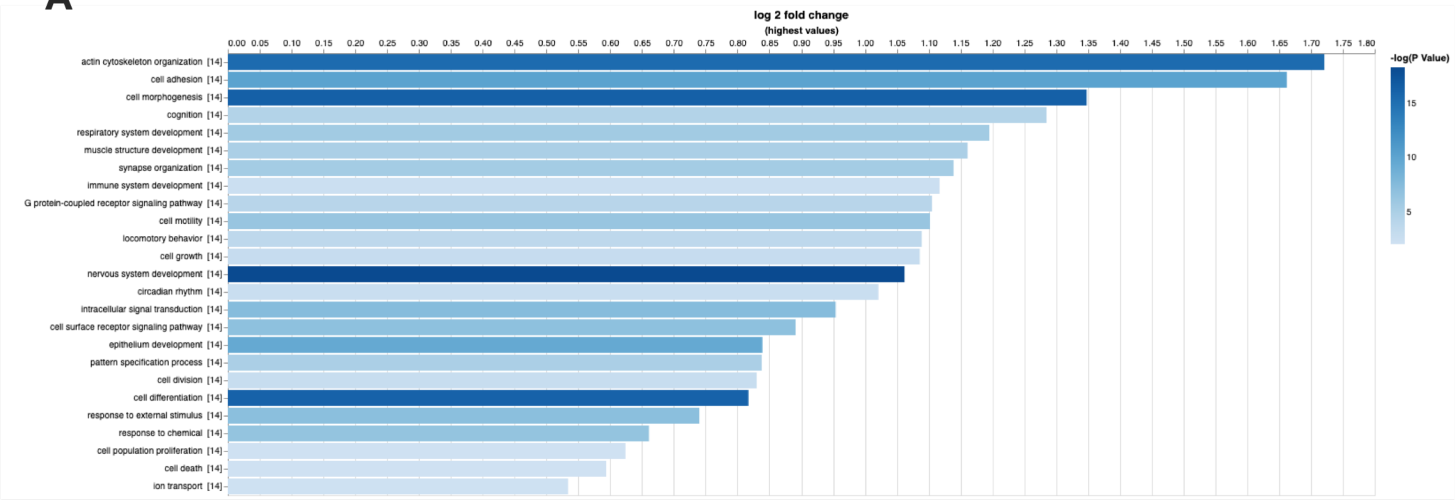

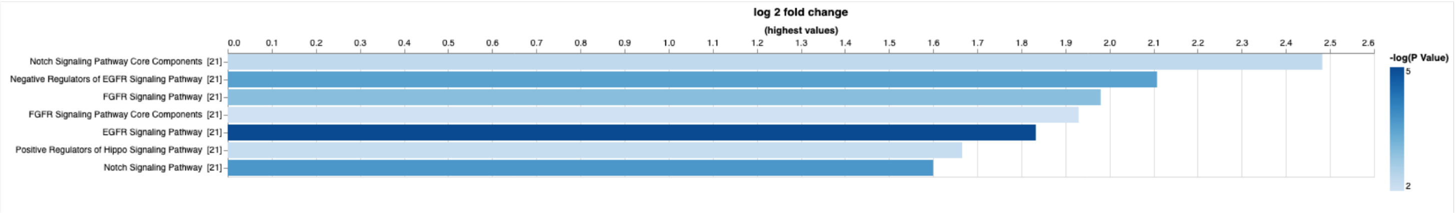

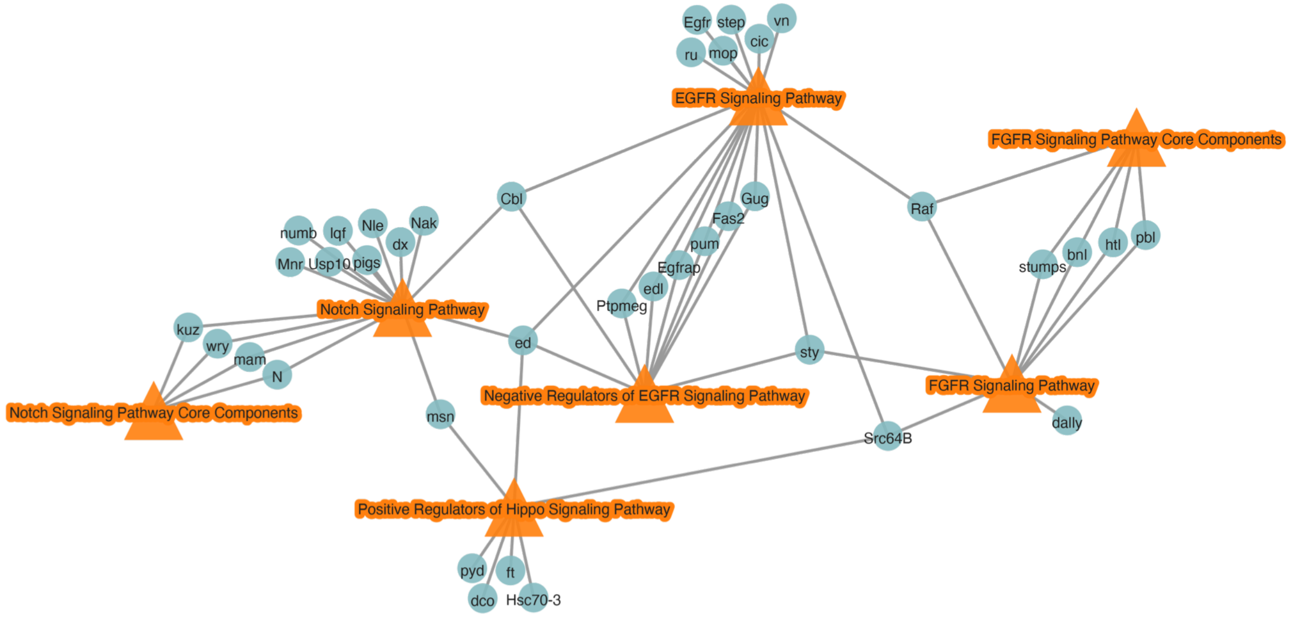

### Supplementary Figure 2

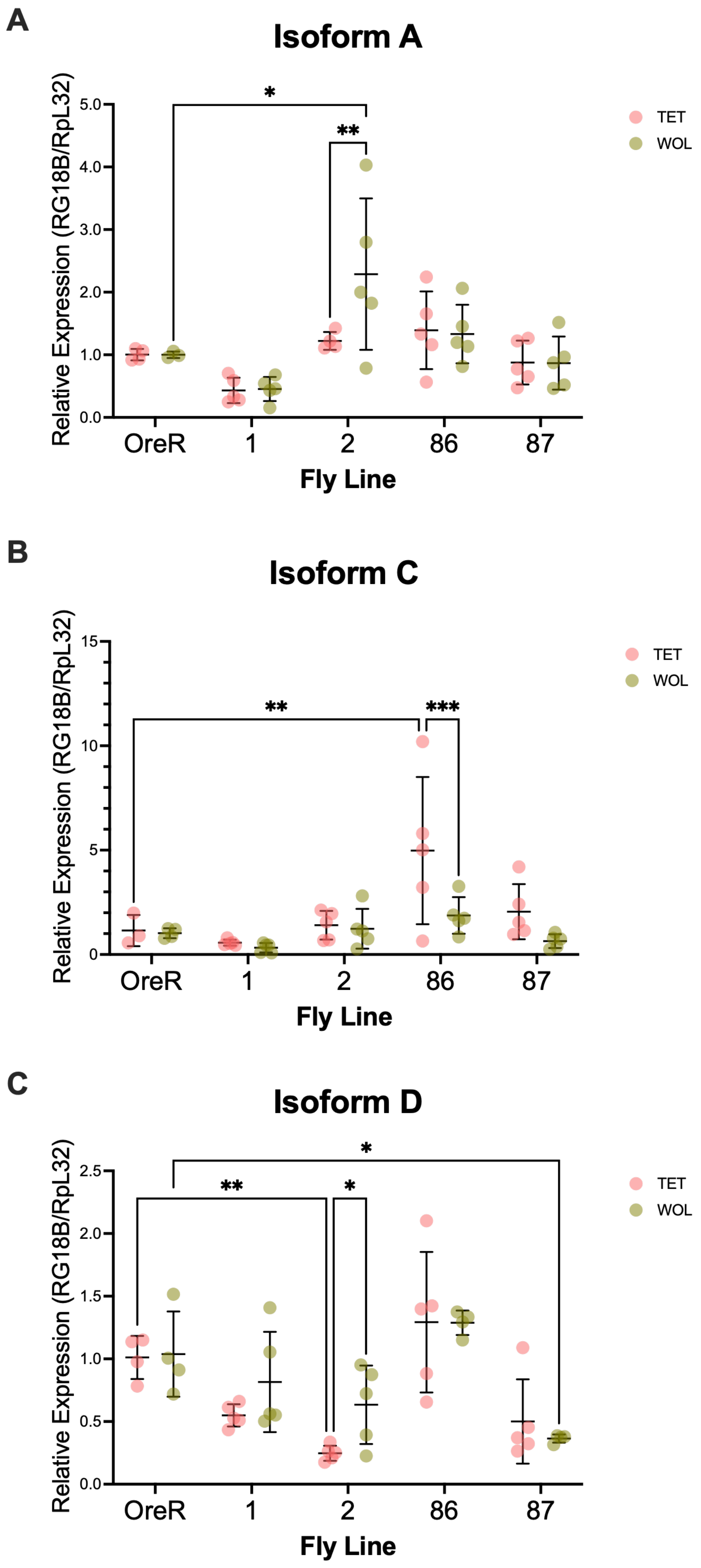
